# Metabolic slowdown as the proximal cause of ageing and death

**DOI:** 10.1101/2023.08.01.551537

**Authors:** J. Wordsworth, P. Yde Nielsen, E. Fielder, S. Chandrasegaran, D. Shanley

**Author notes:** Corresponding authors; &.

## Abstract

Ageing results from the gradual loss of homeostasis, and there are currently many hypotheses for the underlying initial causes, such as molecular damage accumulation. However, few if any theories directly connect comprehensive, underlying biological mechanisms to specific age-related diseases. We recently demonstrated how a specific maintenance system impeding overactivity disorders such as cancer might undergo positive selection while still resulting in a gradual homeostatic shift toward slower metabolism. Here we connect this metabolic slowdown, via a series of unavoidable homeostatic shifts, to the hallmarks of ageing, including mitochondrial dysfunction, insulin resistance (IR), weight gain, basal inflammation, and age-related diseases such as atherosclerosis. We constructed the fuel and energy model (FEM) based on these shifts and found that ageing via metabolic slowdown could explain not only the effects of anti-ageing interventions such as rapamycin and calorie restriction, but many of the paradoxes of ageing that currently defy alternative theories.

## Introduction

Among the many theories describing how and why we age, the disposable soma theory (DST) is the most complete and widely accepted, suggesting the evolutionarily reasons why insufficient somatic maintenance leads to damage accumulation and loss of homeostasis^1, 2^. However, the mechanisms connecting insufficient somatic maintenance as the initial cause to ageing phenotypes and diseases are still the subject of much debate and controversy, with some asking whether the theory is still supported by the literature^3–5^. Others are turning instead to the idea that ageing is a TOR-centric quasi-programme continuing from development that is beyond the power of natural selection to remove^6^. However, with both developmental and damage-based theories there are multiple gaps in the proposed mechanisms that still require elucidation. A recent review of the potential for computational models of ageing highlights this issue by not identifying a single model that demonstrates a connection between initial cause and homeostatic shifts that cause ageing and disease^7^. Other reviews have referred to “ageing-themed models”^8^, detailing many excellent models that look at particular aspects of ageing such as telomere shortening or loss of protein homeostasis. Indeed there are many useful models that offer falsifiable hypotheses for their proposed inducers of ageing such as the recent senescence-based models by Kowald and Kirkwood (2021)^9^ and Karin, et al. (2019)^10^, but in these models cell senescence is both the cause of ageing and the inducer of mortality without reference to the mechanism by which one leads to the other. Although the link between the senescence associated secretory phenotype and the age-associated increase in basal inflammation is implicated, this is far from a comprehensive network that leads to specific homeostatic changes.

In this paper we will show how a different evolutionarily plausible initial cause, manifesting as a single metabolic change, results in homeostatic shifts that induce ageing as represented by the specific hallmarks and diseases observed across all major fauna. The central tenet of this model is that in multicellular, mitotic organisms, mutants are arising constantly within the soma and some will be detrimental to the organism, but not necessarily to the cell. Apoptosis resistance or faster growth, for example, could provide a selective advantage compared to the wildtype cells that allow the mutant to spread.

Current ageing theory has focussed overwhelmingly on mutations that do not impact selection within-tissues, with the suggestion that ageing therefore results in increasing noise. This is undoubtedly occurring and is likely detrimental to tissue function^11^, but offers gaining predictive insight is more challenging as the changes are random. If these mutations were the central cause of ageing, then we could expect age-related deaths to take a wide variety of forms. Cholesterol depletion and cholesterol overload could occur as a result of these stochastic changes. Yet most people die from the same few effectors of mortality, while the change in cholesterol levels is overwhelmingly unidirectional. The prevalence of cancer among these effectors of death is solid evidence that mutants undergoing positive selection are central inducers of homeostatic dysfunction, yet the importance of non-cancerous mutants is only just emerging^12^.

In contrast to noise-inducing mutants, those with a selective advantage will result in non-random changes that predict specific homeostatic shifts in the direction of their selection. Recently, Nelson and Masel (2017)^13^ used computational modelling to argue that in multicellular organisms, these ‘uncooperative’ cells would inevitably takeover proliferative tissues and make ageing an unavoidable consequence of mitotic multicellular somas. However, while their models were highly convincing under the conditions tested, our own models demonstrated that there was a specific mechanism of control that could prevent the spread of uncooperative cells^14^.

We used the example of cells which obtained mutations for accelerated growth, metabolism, and replication (hereon ‘faster’). In the absence of control, these faster cells dominated the tissues as predicted by Nelson and Masel (2017)^13^. However, if cells could compare their growth rate and compete to modify each other in a way that gave an advantage to the slower cells, which we term **selective destruction**, this could not only prevent the spread of faster cells, but could even reverse the selective advantage causing the spread of slower metabolising cells (Figure 1A).

**Figure 1.**
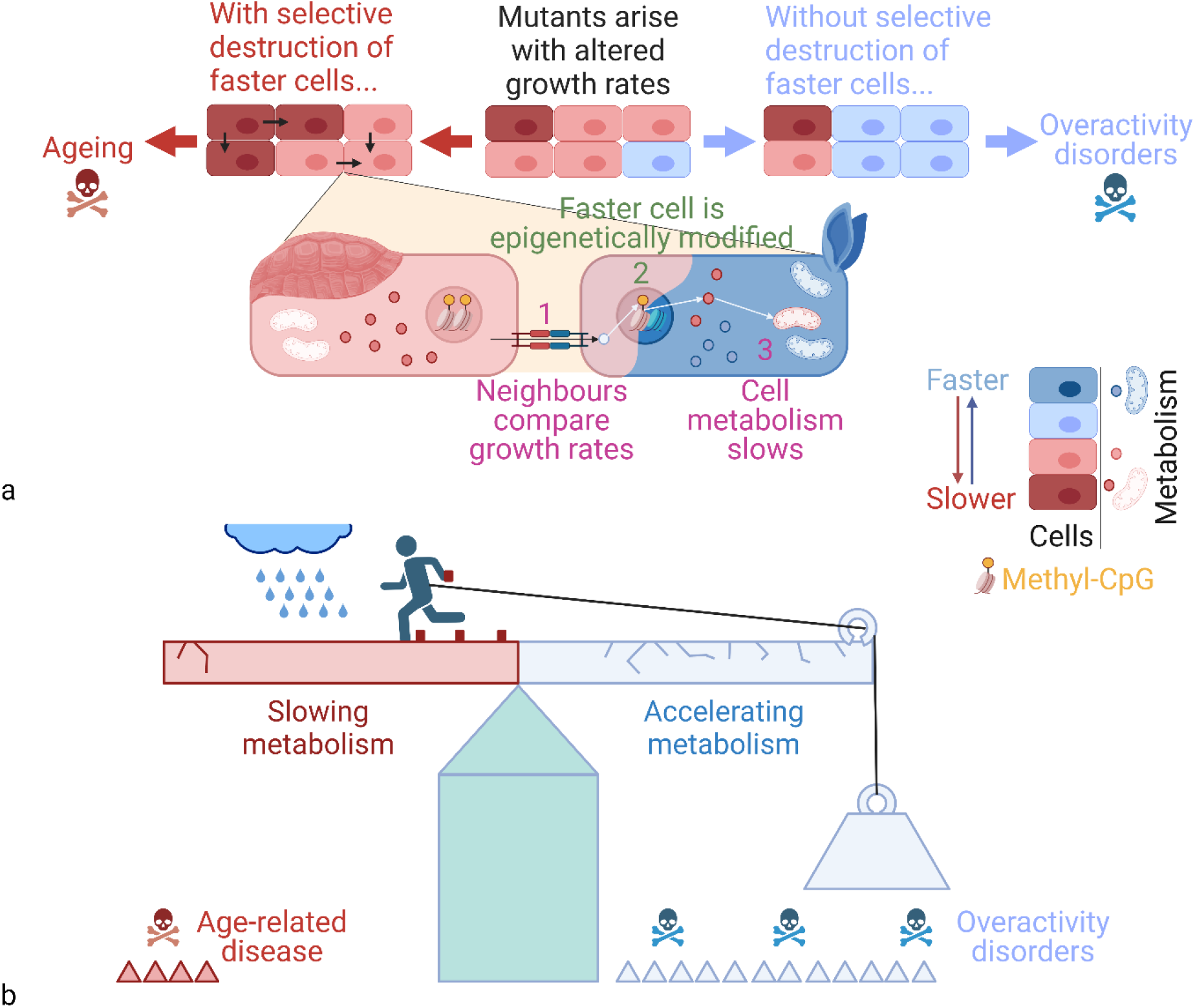
Impact of mutants and surveillance on homeostasis. (A) Fast (blue) and slow (red) mutants occur. Absent surveillance, the faster mutants spread, but with surveillance by selective destruction, the slow mutants spread. (B) The balance between natural fast cell spread (pulled by weight) and selective destruction working to prevent it (running man), but causing spread of slow cells. Cracks show accelerating metabolism is dangerous earlier, rain that insufficient slowing could result in slipping toward accelerating metabolism, and blocks are epigenetic marks preventing slips. Images created with BioRender.com.

While slowing metabolism may initially appear no better than accelerating it, mutants with accelerated growth, metabolism, and reproduction are associated with a host of **overactivity disorders** which make them highly dangerous. Firstly, accelerated replication makes them more mutagenic, leading to increased risk of further detrimental mutations including progression toward cancer. Secondly, they will significantly increase the risk of tissue-specific overactivity disorders. For example, in fibroblasts, which are present in almost all organs, accelerated metabolism will make them more likely to reach the self-sustaining threshold and induce irreversible fibrosis^15^, such as in currently incurable idiopathic pulmonary fibrosis^16, 17^. In other tissues the effects could be equally lethal, such as in β-cells where accelerated metabolism will lead to increased insulin production at the same time as making the cells resistant to apoptosis, with potentially catastrophic effects on glucose homeostasis^18^.

The risks of these overactivity disorders are increased by the presence of even small numbers of faster mutants, whereas even if slower mutants are allowed to spread, they have little detrimental effect, compensated for by the increased activity of wildtype cells. Selective destruction therefore is an active force to prevent the natural dominance and spread of faster cells, resulting in the spread of slow metabolising cells as depicted in Figure 1B. Thus, we suggested that selective destruction of faster cells would be antagonistically pleiotropic, preventing the overactivity disorders which could arise early in life and thus be highly detrimental to fitness, at the cost of a gradual metabolic slowdown. As we argue here, this could be the underlying cause of ageing and age-related disease.

Metabolic slowdown is consistently observed with ageing^19, 20^. In humans basal metabolic rate declines with age independent of the decline in fat-free mass^21–26^. It is also evident in mice^27^, rats^28–30^ and dogs^31^.

In our previous paper, we suggested that the central mechanism of selective destruction is through epigenetic modification, allowing temporary communication between a slow and fast cell to induce semi-permanent changes which slow the faster cell’s metabolism (Figure 1A, lower). We hypothesise these changes are the primary cause of the CpG island (sites with multiple CpGs) methylation at loci corresponding to the epigenetic clocks associated with biological age^32^. Here, we describe how epigenetically slower cells link the evolutionary cause of selective destruction with a comprehensive mechanism of functional decline. This explains not only multiple diseases of ageing and effects of anti-ageing treatments, but also many of the paradoxes which plague other theories of ageing.

### Metabolic slowdown induces mitochondrial dysfunction

Life requires maintaining fuel and energy levels to continue vital processes. Most active processes require energy released from hydrolysing ATP, necessitating tight regulation. Notably, as both ATP synthesis and hydrolysis can be described as ‘metabolism’, we separated **ATP hydrolysing cellular metabolism** (**AHCM**) which utilises ATP for growth and activity, from the **ATP producing mitochondrial metabolism** (**APMM**) which produces ATP through Krebs cycle and oxidative phosphorylation (OXPHOS). **Glycolysis**, which produces ATP but is not mitochondrial is referred to separately, the resulting ATP dynamics shown in equation 1.

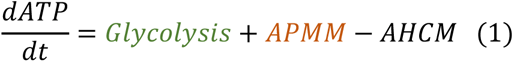

Metabolic slowdown in the context of selective destruction reflects reduced AHCM: cells undergoing less protein synthesis and growing and proliferating more slowly are hydrolysing less ATP. Thus, as AHCM declines either ATP levels will rise, or if ATP is to remain constant, assuming glycolysis is a small fraction of ATP production, then APMM must also decline (AHCM ≈ APMM from equation 1). Consistently, despite decreasing basal and stimulated AHCM with ageing^21, 33^, ATP levels appear remarkably stable. There is little difference in ATP or size of the adenosine phosphate pool in ageing mice^34, 35^. Indeed, muscle-specific Tfam-knockout mice (with reduced mitochondrial gene transcription) have progressively deteriorating respiratory function, but maintain ATP levels by substantially increasing mitochondrial mass^36^. Equally, a study of mononuclear blood cells showed that ATP levels stayed relatively constant in humans from age zero to over 100^37^.

A central regulator of ATP homeostasis is **AMP-activated protein kinase** (**AMPK**) which is activated through an allosteric mechanism by high ratio of AMP:ATP. As such, a key function of AMPK is to restore ATP as levels drop in response to increasing AHCM^38^. Thus, AMPK regulates two main pathways designed to restore ATP levels: firstly, it slows growth and cell cycle by deactivating mammalian Target of rapamycin complex 1 (mTORC1)/mitogen-activated protein kinase (MAPK) pathways, inhibiting AHCM^39^, and secondly it activates transcription factor EB (TFEB) and Peroxisome proliferator-activated receptor-γ coactivator (PGC)-1α, triggering mitochondrial biogenesis and OXPHOS, activating APMM^40^ (Figure 2A).

**Figure 2.**
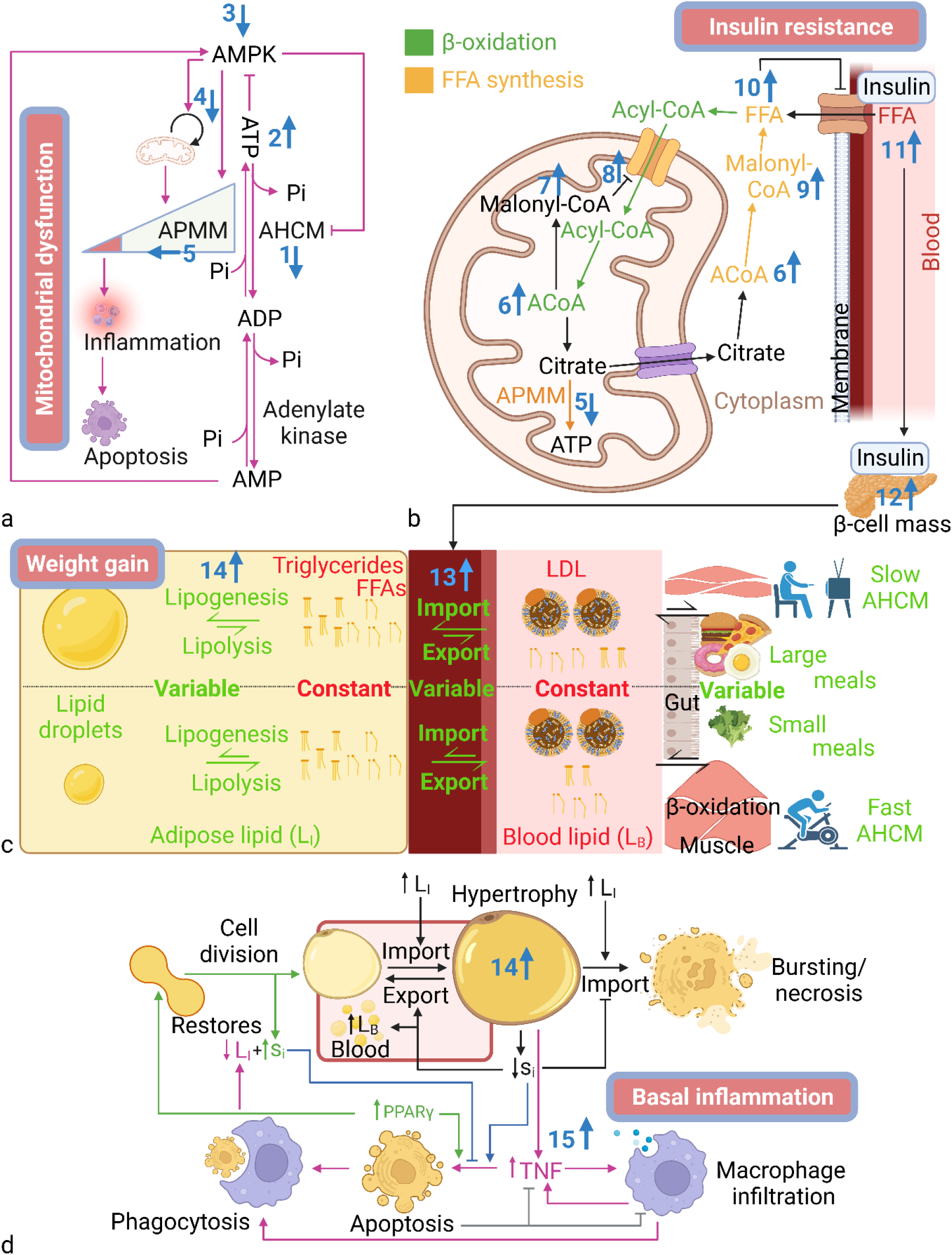
Regulation of energy homeostasis. (A-D) Blue numbers indicate age-related changes and homeostatic effects leading to consequences in blue-bordered boxes. (A) Regulation of ATP dynamics. (B) Detailed model of ACoA dynamics. (C) Regulation of lipid dynamics maintaining homeostasis despite a varying environment. (D) L_I_ homeostasis through inflammation. Images created with BioRender.com. Abbreviations, see table.

Therefore, metabolic slowdown and ATP build-up-induced deactivation of AMPK (by high ATP, low AMP) will lead to reduced mitochondrial ATP production from APMM. The mechanisms by which mitochondrial activity are reduced are highly complex but should include some combination of reduced mitophagy and mitochondrial synthesis, reduced complex activity or production, increased complex degradation, membrane depolarisation, increased ROS production, and thus reduced mtDNA quality. These outcomes could easily be mistaken for the effects of accumulating mitochondrial damage. Indeed, the conventional wisdom is that ROS produced during OXPHOS cause damage accumulation in the mtDNA, reducing mitochondrial function^41, 42^, which then reduces the cell’s capacity for AHCM. Contrarily, we suggest that declining AHCM might be the initial cause, necessitating reduced mitochondrial function.

One aspect of mitochondrial ageing seemingly more consistent with a stress response to accumulating damage is the activation of the inflammasome and initiation of apoptotic pathways in a state described as ‘mitochondrial dysfunction’^43^. However, such a response could also be the desired response to significant metabolic slowdown. When cells become infected, ATP is hijacked for pathogen replication; hence infected cells decelerate ATP production through OXPHOS^44^, activate the immune response, and induce apoptosis. Thus, we speculate that after sufficient metabolic slowdown has reduced APMM below the threshold consistent with infection, it leads to activation of the inflammasome. **‘Mitochondrial dysfunction’ may therefore not reflect dysfunction** at all, but the correct mitochondrial function in response to a perceived state of infection.

If the primary inducer of metabolic slowdown is epigenetic modification, one factor that could distinguish between mitochondrial dysfunction as a homeostatic response and as a result of irreparable damage, is that Yamanaka (OSKM) factor epigenetic rejuvenation can rejuvenate mitochondrial function^45, 46^. If mitochondrial function had declined due to mtDNA damage, the effects should be permanent, not reparable by epigenetic changes, but if mitochondrial function has declined as a homeostatic response this fits with the observed rejuvenation by OSKM. Thus, metabolic slowdown by selective destruction may be the best explanation for three of the central phenotypes of ageing:

- Declining AHCM
- Declining mitochondrial function
- Increased basal inflammation (via mitochondrial dysfunction)

However, mitochondrial dysfunction is merely the first compensatory shift in a long chain.

### Metabolic slowdown induces insulin resistance (IR)

Just as reduced ATP hydrolysis requires a compensatory reduction in ATP production, reduced usage of fuels for ATP production will necessitate its own compensatory shifts to prevent build-up and toxicity. The primary fuels entering the mitochondria are pyruvate from glycolysis and acyl-CoA from the **β-oxidation** of free fatty acids (FFAs). Both pyruvate and acyl-CoA are then converted to acetyl-CoA (ACoA), which is then used for APMM or converted back into FFAs via **FFA synthesis**, as shown in equation 2.

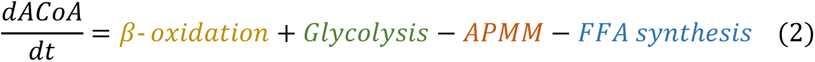

ACoA is regulated by a powerful system of negative feedback through acetyl-CoA carboxylase enzymes 1 & 2 (ACC1 & 2) converting ACoA to malonyl-CoA. In the mitochondria, malonyl-CoA inhibits the transport of further acyl-CoA into the mitochondria (inhibiting β-oxidation), while in the cytoplasm malonyl-CoA is the substrate for fatty acid synthase, which converts it to FFAs (Figure 2B). ACoA can thus be kept relatively constant by balancing rates of APMM with corresponding changes to β-oxidation and FFA synthesis. Thus, as APMM declines and less ACoA is passed into Krebs cycle for ATP synthesis, the excess ACoA is converted to malonyl-CoA which both inhibits β-oxidation and activates FFA synthesis, reducing ACoA.

Notably, FFA synthesis is limited mainly to the adipose and lipogenic tissues, so most organs will adapt to lower APMM primarily by inhibiting β-oxidation. However, both changes have the same consequence of increased intracellular FFAs.

In non-adipose tissues where lipids cannot be easily stored, excess FFAs quickly become toxic to metabolism^47–49^, and must be kept close to equilibrium by reducing FFA import into cells. **Import** is controlled primarily by **insulin** through the activation of cluster of differentiation 36 (CD36)^50–52^. Non-adipose tissues with excess FFAs therefore reduce their **insulin sensitivity** (**s_i_**), such as by reducing CD36 expression, reducing their import to match their slower metabolism (APMM and AHCM), preventing lipid overload. Thus, we observe that **metabolic slowdown inevitably leads to IR**, which is one of the key hallmarks of ageing, highly associated with mortality^53^.

### Metabolic slowdown induces weight gain

Following our chain of causation, removing less FFAs from the blood will (without compensation) result in increased FFAs in the blood. Unlike glucose, FFAs are not water soluble, so are mainly transported as esters on lipoproteins. For simplicity, the combined **blood lipids** are referred to as **L_B_**.

Notably, L_B_ undergoes significant changes during ageing^54, 55^ in stark contrast to blood glucose. However, there are still potent control mechanisms designed to restore L_B_ equilibrium, even if they are not as strong as for ATP, likely reflecting a ‘healthy range’.

Although the non-adipose tissues with excess FFAs are becoming increasingly insulin resistant, the adipose tissue has an additional mechanism to prevent the toxicity of intracellular FFAs by packaging up the esters (triglycerides) as lipid droplets (LDs) which are coated in cholesterol-rich, hydrophilic membranes protecting the adipocytes from the insoluble molecules^56^. Thus, adipose can store fat, maintaining L_B_ homeostasis by regulating adipose import and export according to changes in L_B_, such as from **meals** (equation 3).

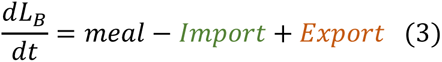

As L_B_ increases, the β-cells in the pancreas produce additional insulin, which increases adipose import and reduces export. By co-regulating import and **export** with lipogenesis and lipolysis which convert FFAs to LDs and vice versa, the adipose can maintain **intracellular lipids** (**L_I_**) like triglycerides and FFAs in all tissues (including itself) as well as L_B_, while changing only the level of stored LDs (still here referred to as L_I_) in response to metabolic or dietary change (Figure 2C).

Thus, the **proximal effects of metabolic slowdown are likely to be weight gain and IR** as we store excess lipids in LDs. However, as metabolic slowdown continues, the adipose cannot provide an infinite store for excess FFAs.

### Metabolic slowdown induces basal inflammation

There is little evidence for a tightly regulated equilibrium value of L_I_ such as a maximum weight per individual^57^. However, one excess calorie absorbed does not equal one excess calorie stored, suggesting we do have mechanisms to curb increasing L_I_, and these become increasingly active as the energy balance becomes increasingly positive^58, 59^.

Notably, although adipose is a proliferative tissue, with around 10% turnover per year in humans^60^, most studies suggest that the number of adipocytes in adulthood remains relatively fixed (with equal rates of death and proliferation)^60–62^. Instead, the adipose mass grows and shrinks mainly by changing cell size^63^, suggesting that L_I_ storage will be finite. Although, the adipose is uniquely capable of avoiding IR by storing its L_I_ in non-toxic lipid droplets; eventually increasing storage results in adipocyte hypertrophy, upon which continuing import leads to bursting and necrosis (i.e. uncontrolled cell death), reflecting mainly stress conditions rather than normal adipose homeostasis^64, 65^. Therefore, to avoid bursting, adipocytes regulate import of L_B_ with respect to their L_I_, firstly by reducing insulin sensitivity (s_i_). However, as in non-adipose, preventing FFA import will increase L_B_, contrary to adipose function. We therefore predicted an additional mechanism to lower L_I_.

Although the cause of inflammation in obesity is often stated as “unknown”^66^, most evidence suggests it plays a central role in the induction of **apoptosis** in hypertrophic adipocytes and their subsequent phagocytosis by macrophages. Obesity is associated with an increase in macrophage infiltration^67^, and inflammatory M1 polarisation^68^. Additionally, immune-deficient mice have increased fat mass^69, 70^ and immune reconstitution of immunodeficient mice on high-calorie diets reduced weight gain^70^.

In both lean and obese mice, Cinti, et al. (2005)^71^ found that more than 90% of macrophages localized around large or dead adipocytes forming crown-like structures (CLS), suggesting fat removal by apoptosis and ingestion of hypertrophic cells. Therefore, L_I_ dynamics are shown in equation 4.

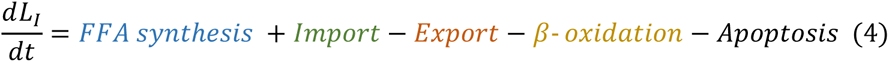

As metabolic slowdown increases FFA synthesis and import, and reduces β-oxidation, the increasing L_I_ requires increased apoptosis to remove the hypertrophic cells, which requires increased basal inflammation.

In adipose, the main apoptosis pathway appears to be extrinsic, triggered by resident macrophages which secrete factors promoting apoptosis^65^. Thus, as indicated in Figure 2D (pink), hypertrophic adipocytes secrete cytokines such as TNF-α which attract macrophages, which in turn produce cytokines to trigger the apoptosis of adipocytes.

Intriguingly, one study using the FAT-ATTAC model to induce adipocyte apoptosis found that apoptosis itself recruited M2 anti-inflammatory macrophages, while living adipocytes were required to induce metabolic switching to pro-inflammatory M1 macrophages^72^. Consistently, Haase, et al. (2014)^73^ found macrophages mainly proliferate at CLS, where they are inducing apoptosis, but proliferation produces mainly M2 macrophages, presumably providing the negative feedback that inhibits inflammation after the hypertrophic adipocytes are removed (Figure 2D, grey). These results indicate that despite the appearance that inflammation sustains itself in a counterproductive way, as suggested by damage-centric and inflammaging models, the pro-inflammatory state is retained mainly by hypertrophic adipocytes as a necessary homeostatic pathway.

As shown in Figure 2D (green), this mechanism returns both L_I_ and s_i_ to equilibrium levels because PPARγ activates both apoptosis and adipocyte differentiation from stem cells^74^, replacing L_I_-laden, insulin resistant hypertrophic cells with smaller insulin sensitive ones, also explaining why adipocyte number is constant^60^. Crucially, inflammation-deficient transgenic mice had reduced rates of adipocyte proliferation and fewer total adipocytes than wildtype mice, which was associated with hepatic steatosis and metabolic dysfunction^75^.

Therefore, the predicted effects of age-related metabolic slowdown would be weight gain^76^, increasing basal inflammation^77^, increased IR^78^, and slower mitochondria^79, 80^, which are among the clearest phenotypes of ageing. Notably, ageing-induced IR was dependent on changes in body composition^78^, which is also consistent.

There is some evidence that these shifts in themselves might be responsible for functional decline and disease. For example, in bone, IR promotes stem cell differentiation to osteoclasts while inhibiting osteoblast differentiation^81, 82^, which results in increased degradation of bone tissue, potentially contributing to osteoporosis. Equally, IR in osteoblasts reduces osteoblastic lifespan^83^, while the increased inflammatory cytokines, particularly IL-1, IL-6, and TNF-α, form a bone resorptive milieu^84^. Thus, while osteoporosis appears like wear-and-tear, it could simply reflect homeostatic shifts in response to metabolic slowdown.

Notably, while subcutaneous fat is often associated with healthy metabolism and insulin sensitivity, visceral fat is often associated with IR, morbidity, and mortality^85^. As basal inflammation reduces rather than inhibits increasing L_I_, eventually metabolic slowdown will lead to the exhaustion of subcutaneous adipose stores. Potentially, the enlargement of visceral adipose reflects the body’s response to this state. These adipose stores will therefore be associated with IR and inflammation, not because of any qualitative metabolic differences, but because they are an emergency depot to remove excess lipid. A further factor is the proximity of visceral fat to important organs, spreading basal inflammation where its effects are less positive.

However, even as our muscle and other tissues are replaced by adipose, the rising IR is preventing tissues from accepting L_B_, provoking levels to rise. Declining appetite with age is likely one compensatory mechanism^86^, reducing lipid absorption from the gut. We also have a process of reverse-cholesterol transport (RCT), which allows us to remove excess lipid and lipoprotein in the bile. However, while RCT will provide a buffer against increasing L_B_, its capacity will also be finite, dependent on the number and capacity of liver cells to produce bile, likely also reduced by declining AHCM in a similar way to adipose storage. Eventually, even if L_B_ is maintained at relatively robust equilibrium, it will increase because of metabolic slowdown.

### Metabolic slowdown induces atherosclerosis

While rising L_I_ induces little detriment outside increasing basal inflammation, rising L_B_ can have serious effects on health. It is well established that one of the proximal events in atherosclerosis is the deposition of lipid in the blood vessel walls^87, 88^. The damage-centric hypothesis suggests this is a “response to injury” in the vessel wall^89^. “Chronic low-level insult” leads to endothelial dysfunction and lipid accumulation^87, 90^. Alternatively, we suggest that a potential cause might be as simple as the loss of L_B_ homeostasis. Rising L_B_ increases the pressure to import the excess to the tissues, while the tissues are increasingly (insulin) resistant to accepting it. Sandwiched between these opposing forces, soluble lipids such as cholesterol can recrystalise out of solution^91^, while insoluble lipids can be deposited in the intima of blood vessels, forming the initial stages of atherosclerotic plaques.

Observations that endothelial dysfunction is the proximal event in atherosclerosis^92, 93^ are based primarily on the location of plaques at dysfunctional sites, but just as an overburdened pipe bursts at the weakest point, L_B_ is likely deposited at these sites because they are the most susceptible sites for accepting it, which does not mean their existence is the primary cause. Of course, increasing endothelial dysfunction likely shifts the equilibrium so that there is more plaque growth at lower L_B_, just as weakening a pipe will accelerate the point at which it bursts.

It has been suggested that low density lipoprotein (LDL) itself is not atherogenic because after macrophages intake LDL they downregulate LDL-receptors (LDL-Rs) until the lipids have been degraded, thus preventing lipid overload and foam cell formation^94^, which is the next major step in atherogenesis. However, oxidised LDL upregulates the macrophage LDL-R^95^, thus causing the lipid overloaded foam cells to become embedded in the growing plaque. This is widely interpreted as evidence that damage accumulation from ROS is the causal factor behind atherosclerosis^94^, but could instead result from higher basal inflammation (hence increased oxidation) combined with the presence of excess LDL due to metabolic slowdown. Consistently, overexpression of the macrophage-expressed enzyme 15-lipoxygenase which promotes LDL oxidation^96^ *protected* rabbits from atherosclerosis^97^, which makes sense if LDL-oxidation is instigated by macrophages in response to quantities of LDL that cannot be removed without inhibiting LDL-R downregulation. By this hypothesis, foam cell formation and atherosclerosis occur not because of excess LDL oxidation, but because LDL oxidation is insufficient to remove the excess LDL and return L_B_ homeostasis. **Atherosclerosis is thus the inevitable result of metabolic slowdown**.

### Metabolic slowdown is a proximal cause of ageing and death

Here, we are proposing that ageing results from epigenetic changes designed to slow down cells in tissues which would otherwise tend to accelerate metabolism over time, which we have termed selective destruction theory (SDT). A chain of causation leads to mitochondrial dysfunction, IR, weight gain, basal inflammation, and finally atherosclerosis, one of the primary inducers of age-related mortality^98^, as summarised in Figure 3A. Notably, Alzheimer’s and various types of neurodegeneration have been linked to mitochondrial dysfunction^99, 100^, IR^101^, dyslipidaemia^102^, inflammation^103^, and a decrease in cerebral blood flow^104^, which may reflect in part the atherosclerotic conditions of cerebral blood vessels^105, 106^. However, it should be noted that declining AHCM itself could be responsible for much of the age-related dysfunction.

**Figure 3.**
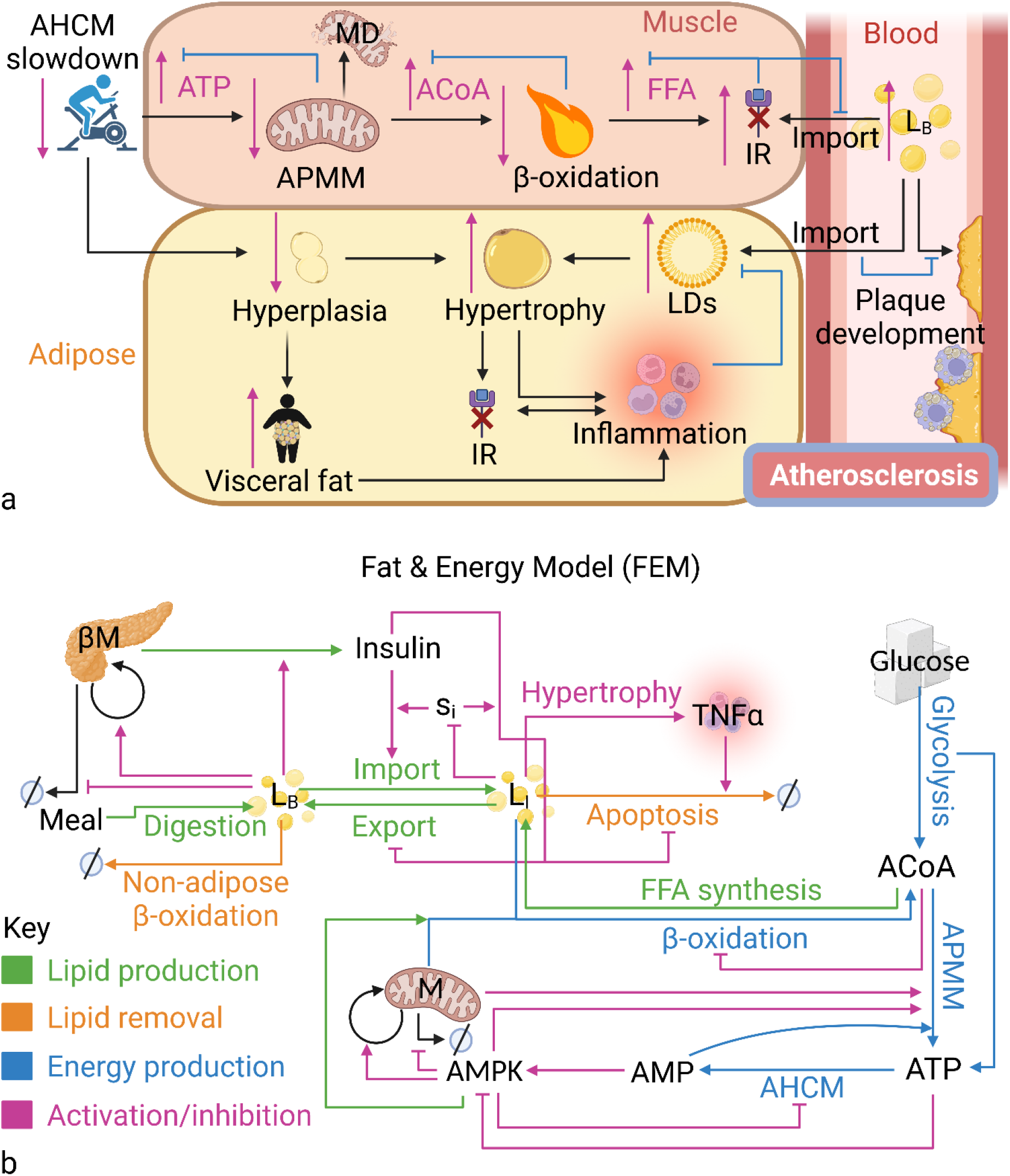
Fuel and energy homeostasis and model. (A) Knock-on effects of declining AHCM that result in increasing L_B_ and risk of atherosclerosis. (B) Schematic of the Fuel & Energy Model (FEM). Images created with BioRender.com. Abbreviations: see table.

For example, primary sarcopenia (loss of muscle mass) is central to age-associated frailty^107^, as is the mass-independent loss of strength, termed dynapenia^108^. Both could result directly from declining AHCM as reflected by the 2-3% decline in basal metabolic rate in muscle every decade after age 20, and 4% after age 50^33^. The gradual degradation of our extracellular matrix (ECM) with age^109^ could reflect a decline in the fibrotic capacity of fibroblasts with slowed protein synthesis. In a fascinating modelling study, Karin and Alon (2017)^18^ showed how decreased cell proliferation with age might predispose individuals to type II diabetes.

Additionally, our reduced stress responses including the antioxidant^110^ and DNA damage response^111–113^ which are often touted as evidence of ageing by damage accumulation^114^, could simply be the consequence of reduced TOR-driven metabolism such as protein translation. Indeed, it has been suggested that nucleotide excision repair declines as a result of reduced levels of the associated proteins^115^. Failure of the unfolded protein response (UPR) with age^116^ is also likely contributed to by endoplasmic reticulum stress due to higher lipid levels.

To test the predictions of SDT we created the Fuel and Energy Model (FEM) based on mathematical representations of equations 1-4 and equations S1-4 defined and justified in the supplementary methods according to the evidence above. Briefly, AHCM is a function of the basal rate of cellular metabolism (k_CM_) and environmental elevation (eg. exercise), inhibited by AMPK. APMM is activated by AMPK and intrinsic mitochondrial capacity (IMC), the latter being a function of mitochondrial mass and functionality, with AMPK stimulating IMC increase and inhibiting decrease. AMPK activity is determined by the ratio of AMP:ATP. L_I_ is converted to ACoA by β-oxidation, also dependent on IMC and AMPK, and inhibited by ACoA, which is the substrate for FFA synthesis, regenerating L_I_. L_I_ can either be exported to the blood, forming L_B_, or removed when hypertrophic cells undergo apoptosis. Apoptosis is stimulated by L_I_ and inflammation, the latter modelled as a single cytokine, TNF-α, which increases with L_I_ (reflecting additional hypertrophy). Adipocyte apoptosis is inhibited by insulin (induced survival signalling). Insulin is produced by β-cells, dependent on the **β-cell mass** (**βM**), which is a function of β-cell proliferation and apoptosis, stimulated and inhibited by L_B_, respectively. Insulin stimulates L_B_ import, forming L_I_, just as it inhibits export and apoptosis all dependent on the insulin sensitivity (s_i_) of the cells, which is decreased by L_I_ (reflecting hypertrophy). The model schematic is shown in Figure 3B. Detailed outputs of the model are shown in the supplementary results (Figures S1-5).

When we steadily reduced the basal AHCM rate (k_CM_, Figure 4A), causing total AHCM to decline (not shown), the FEM maintained ATP (Figure 4B) by reducing IMC (not shown), predicting decreasing mitochondrial function, which reduced APMM (Figure 4C). ACoA was maintained by corresponding decrease in β-oxidation (Figure 4D-E), which shifted L_I_ to a higher equilibrium (Figure 4F), reducing s_i_ (Figure 4G) predicting weight gain and IR, and increasing inflammation and apoptosis to retard the L_I_ increase (Figure 4H-I), predicting basal inflammation. Notably, the FEM did not predict atherosclerosis as L_B_ was kept in equilibrium by increasing βM and insulin levels (Figure 4J-L). However, when we introduced a maximum βM (see supplementary model discussion and Figure S6 for further detail), reflecting that the pancreatic islets cannot expand forever, once βM plateaued, rising insulin could no longer compensate for additional lipid, and L_B_ began to rise (Figure 4J-L), thus predicting atherosclerosis.

**Figure 4.**
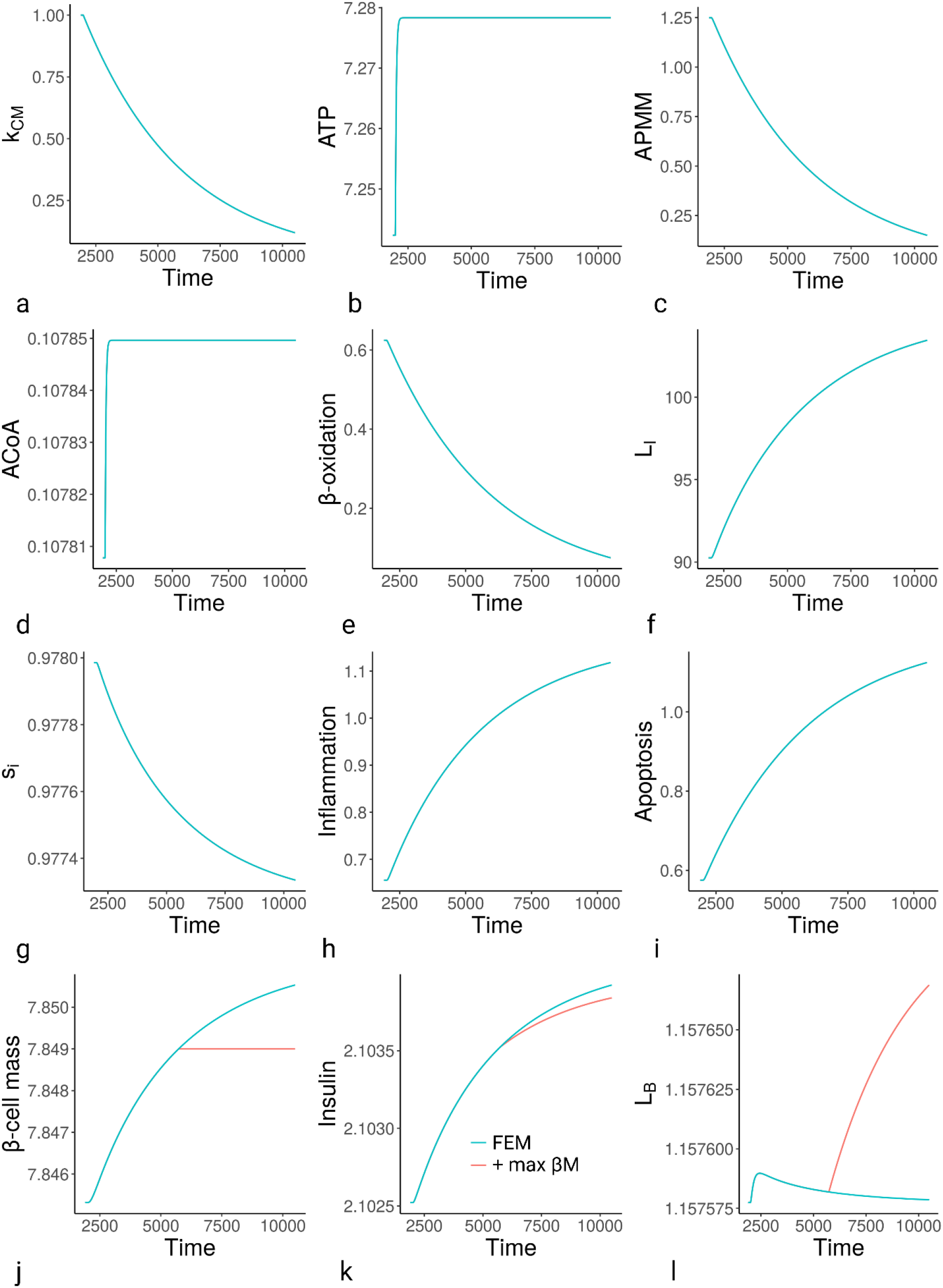
Simulations of the FEM in response to declining k_CM_. (A) k_CM_ input. (B) ATP. (C) APMM. (D) ACoA. (E) β-oxidation. (F) L_I_. (G) s_i_. (H) Inflammation. (I) Apoptosis. (J) β-cell mass. (K) Insulin. (L) L_B_. Simulations start at t=1900 au, and k_CM_ declines according to supplementary equation S5 from t=2000 au. All graphs include effects of a maximum βM of 7.849 au as red line; however, it only affects Figures J-L. Abbreviations, see table.

We suggest that this is the most comprehensive and detailed model of ageing currently available, particularly outside the pathways proposed for ageing via damage accumulation. We therefore tested how the FEM would respond to anti-ageing interventions.

### Anti-ageing interventions and metabolic slowdown

We modelled calorie restriction (CR) in the FEM as a reduction in the meal value shown in equation 3. As CR did nothing to prevent the underlying metabolic slowdown, not affecting ATP or AHCM (Figure S2P & W), its beneficial effects occurred by reducing L_I_ and basal inflammation and increasing s_i_ (Figure S2S & U-V), consistent with the literature^117, 118^.

To observe how CR would impact mortality, we utilised rising L_B_ as a proxy for atherosclerotic risk. When we swapped CR to ad libitum (AL), all variables including L_B_ (Figure 4A) returned to the levels of continuous AL. Importantly, Mair, et al. (2003)^119^ swapped flies between dietary restriction (DR) and AL and found mortality profiles that matched closely to those produced by the FEM (see their Figure 1B for comparison). These profiles suggested that CR was primarily risk-preventative, affecting only the ultimate inducer of atherosclerosis by reducing L_B_, but not affecting the underlying cause of slow L_B_ accumulation (ie. selective destruction and metabolic slowdown). Notably, the authors suggested the findings were inconsistent with CR affecting ageing via reducing damage accumulation as such interventions should have lasting effects (as less damage has accumulated at the point the intervention ceases).

When the same lab tested swapping between CR and AL in mice, they observed some memory effect^120^, suggesting CR may have gero-protective properties in mice, if not flies. Notably, multiple studies and meta-analyses indicate the beneficial effects of exercise on dislipidemia and mortality, and the benefits are not limited to the exact times the individuals are exercising^121, 122^, but are limited to healthspan and mean lifespan with no maximum lifespan extension^123–125^. These long-lasting benefits may thus still be primarily risk preventative rather than gero-protective. In asking whether the FEM could also predict this, we considered a more realistic risk assessment would reflect the levels of atherosclerotic plaques from excess L_B_. While L_B_ can change quickly, plaques are both formed and degraded more slowly, low L_B_ allowing net degradation, and higher L_B_ shifting the parameters so plaque formation is faster than degradation (see equation S3.10). From this we modelled regimens of CR (low meal) and exercise as temporary increases in AHCM, demonstrating that both regimens could induce long-lasting benefits while still being entirely risk preventative (Figure 4B). However, these results do not rule out additional gero-protective effects.

The most obvious predictor of gero-protective effects, at least for SDT, is via the epigenetic clocks capable of predicting biological age by the quantity of specific CpG island methylation. The exact purpose of these CpG methylations is unknown, and has led some to suggest that their clock-like appearance is evidence of programmatic ageing^126^. However, if SDT is correct, these CpG methylations reflect the outcomes of cell communication in which slow cells induce reprogramming of faster cells by permanently slowing their AHCM. Although speculative, if true, these epigenetic marks will be the truest biomarkers of ageing, and the best evidence of a gero-protective treatment outside extended maximum lifespan (and be largely independent of DNA damage). Notably, CR^127^ does affect the epigenetic clock, and extends maximum lifespan in mice^128, 129^.

Metformin is primarily regarded as an AMPK activator, but its mechanisms are not well understood^130^. We addressed this in the FEM with changes to several parameters (see supplementary methods), impacting AMPK. When metformin was modelled to affect rates of AMPK activation and deactivation, it had little impact on L_B_ or plaques (Figure S5B-C), but increasing the impact of AMPK on mitochondrial activity (IMC, Figure 5C and S5-E-G) resulted in robust L_B_ and plaque reduction (Figure 5D-E), predicting that metformin has risk-preventative effects. Notably, there is little evidence that metformin extends maximum lifespan in mice^131^ or flies^132^ and though it extends maximum lifespan in *C. elegans*, this is reversed when the gut microbiome is removed, suggesting it is the effect on bacteria which extends lifespan^133^. Metformin also failed to affect the rate of CpG methylation, suggesting it is not geroprotective, while the mTORC1 inhibitor rapamycin reduced CpG methylation and the estimate of biological age^134^.

**Figure 5.**
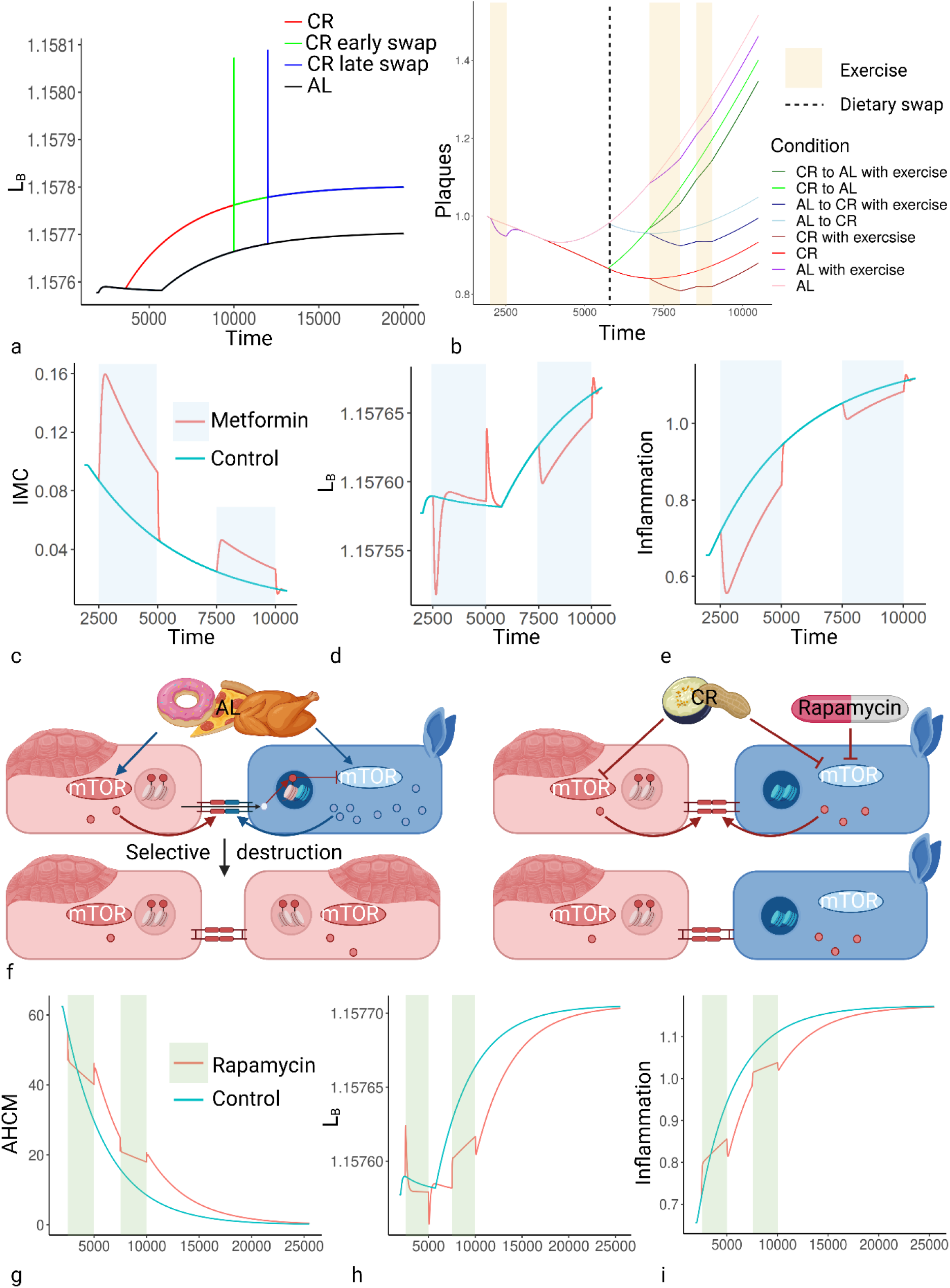
Effects of anti-ageing interventions on energy homeostasis. (A) Effects of reduced Meal (mimicing CR) on L_B_. (B) Combined effects of CR and exercise regimens on mortality as a function of plaque number and size. (C-E) Effects of metformin (as represented by a reduction in λ_4_, see equation S1.9 and Figure S5E-G) on: (C) ATP; (D) inflammation; (E) L_B_. (F) Impact of CR and rapamycin on mTOR signalling and selective destruction. (G-I) Effects of rapamycin (represented by changing k_CM_, see Figure S5H-M) on: (G) AHCM; (H) L_B_; (I) Inflammation. Images created with BioRender.com. Abbreviations, see table.

Rapamycin is probably the most consistent longevity enhancing drug yet documented, with several studies showing it affects maximum lifespan in mice^135, 136^ and flies^137^. Consistently, rapamycin has long-lasting effects on mortality even after the regimen ceases^138^. It has multiple effects on cell and molecular phenotype, including reducing cell division rate, but these are all (at least at low dose) thought to result from the inhibition of mTORC1^139, 140^, which is a central driver of cell growth and metabolism^141^, essentially driving AHCM. Fascinatingly, this is essentially mimicking metabolic slowdown, which the FEM predicts should accelerate ageing, the increase of L_I_, basal inflammation, and atherosclerosis. Notably, rapamycin has been shown to induce IR, glucose intolerance, and diabetes, while still being associated with robust increases in lifespan^142, 143^, suggesting it can have the negative effects predicted by the FEM, but may be having other compensatory effects, such as reducing the rate of selective destruction.

By slowing the rate of cell growth/division, rapamycin may reduce either mutation rate or **increase the growth similarity between fast and slow cells** as shown in Figure 5F, by reducing the rate of fast cell growth toward the level of slower cells, and therefore reducing the rate of selective destruction. To demonstrate the effects of this, we modelled rapamycin to both cause a reduction in AHCM and decelerate the rate of metabolic slowdown (Figure 5G), which would be the primary effect of reduced selective destruction. We compared rapamycin to untreated controls, and consistent with the literature^138^, rapamycin had long lasting beneficial effects on L_B_ and basal inflammation (Figure 5G-I), as well many other parameters (Figure S5F-M). However, it also showed that taking rapamycin at the wrong time could cause L_B_ to spike (Figure 5H) and exacerbate basal inflammation temporarily (Figure 5I), suggesting the drop in AHCM could have detrimental consequences in the short term. Consistently, rapamycin has transient (paradoxical^144^) negative effects on weight gain and fasting glucose levels at the start of treatment^145^. Schindler, et al. (2014)^146^ found that key factors affecting metabolic impairment from rapamycin included the dose and the age of treatment, which both predict the benefit-cost ratio shown in Figure 5F-I. Other explanations, such as the mTORC2-specific rapamycin effects^143^ do not explain the general paradox of growth signalling and ageing^147, 148^. Insulin-like growth factor (IGF) and growth hormone (GH) show robust decreases during lifespan^149, 150^, GH level correlates negatively with cardiovascular risk^151^, and adult supplementation has had short-term beneficial effects^152^, but longer-term use in healthy older adults is associated with increased adverse events^153^, and knockouts are associated with increased longevity^154–156^. The FEM and SDT predict exactly these effects: the short-term benefits of GH supplementation reflect the accelerated metabolism, temporarily reversing the effects of ageing, but encouraging rapid growth and metabolism will only increase selective destruction by making the faster cells most responsive to these stimuli appear more aberrant, thus promoting more rapid ageing. Conversely, knockout mutants may have initially slower metabolism, but the molecules knocked out are the key architects of differences in growth rate between cells, decelerating selective destruction and metabolic slowdown (Figure 5F). Notably, the same reduction of cell growth rate is true for CR, suggesting why it retards the epigenetic clock and increases maximum lifespan; but while rapamycin inhibits mTOR in spite of high nutrients, which results in IR and problems maintaining metabolite levels, CR modulates mTOR by removing the activating nutrient signals, thus maintaining insulin sensitivity.

It should be noted that the FEM was not built to include targets of anti-ageing interventions. The fact that these can be input so easily, and the model responds so realistically, is good evidence that the suggested mechanisms are relevant to ageing.

Finally, the Yamanaka factors and parabiosis have both shown robust life extending potential^157, 158^, indicating epigenetics as an anti-ageing target. Selective destruction induces semi-permanent epigenetic slowing, but we did not predict this would be unidirectional^14^.The *advantage* to slower cells would not be total, with faster cells capable of accelerating AHCM in line with the presumed wildtype (i.e. by comparison to multiple cells). Cells could lose the epigenetic markers slowing metabolism, essentially rejuvenating them. This may partly explain why the exposure to young blood cells rejuvenates old mice, although selective destruction in blood might look quite different to solid tissues. Yamanaka factor rejuvenation essentially removes all epigenetic alterations determining both cell fate and metabolism, returning the cell to the default state. SDT suggests that once the safety of this tool has been improved, it will provide the most effective rejuvenation. However, the effects will not be wholly positive.

### Negligible senescence, ageing, and cancer in light of selective destruction

As we have mentioned above, one of the primary reasons for selective destruction and metabolic slowdown are to prevent the overactivity disorders such as cancer and fibrosis, which notably increase with age. This may be explained partly as increasing basal inflammation has also been associated with both pro-tumorigenic^159^ and pro-fibrotic microenvironments^160^, increasing risk of these disorders with age, ironically induced by selective destruction. Increasing IR may also be pro-fibrotic^161^. Additionally, while ageing may or may not reflect accumulating damage, cancer most definitely does. As such, the risks will increase with age as damage accumulates. In evidence that metabolic slowdown reduces cancer risk, there is a drop in cancer risk in the oldest old^162^, reflecting slower cell division^163^, and cancer risk is notably inverse with Alzheimer’s disease^164, 165^, consistent with the idea the selective destruction is causing one and preventing the other.

CR and rapamycin show little evidence of increasing risks of cancer or fibrosis, which appear to be decreased by the interventions^166, 167^; however, neither intervention targets selective destruction or the epigenetic markings *per se*, instead reducing the rate of cell metabolism, which reduces both the risk of overactivity disorders and (the requirement for) selective destruction. As a result, these treatments can be wholly beneficial but can only delay ageing. Epigenetic rejuvenation such as via the Yamanaka factors and parabiosis could potentially reverse ageing, but will also remove the controls on faster mutants, increasing the risk of overactivity disorders. Thus, ‘negligible senescence’ as we understand it, may be incredibly difficult to achieve. We would merely be trading metabolic slowdown for metabolic acceleration and the associated risks. Notably, the relative abundance of ageing in wild organisms combined with the near absence of negligible senescence is good evidence that evolving lifespan extension is highly problematic. This is contrary to damage-centric hypotheses, where negligible senescence should be achievable by the sufficient upregulation of maintenance.

Disposable soma theory explains this absence by suggesting that no animal in the vast array of niches gains sufficient benefit from this upregulation, because all niches will advantage other energy intensive processes more. SDT suggests that reducing selective destruction and decelerating metabolic slowdown will induce alternative mortality risk. A similar antagonistic pleiotropy has been previously suggested for the evolution of the cancer-protective but pro-ageing p53^168, 169^. SDT expands on this evolutionary plausibility with a comprehensive, modellable ageing process, but as with all antagonistic pleiotropy, the proposed trade-off must survive empirical testing to ensure the cost and benefit are unavoidably linked, and the formal evolutionary substantiation or invalidation of SDT is yet to come.

The study of ageing has largely drifted away from attempts to understand the underlying cause. The inconsistency of our existing damage-based theories with evidence that damage accumulation frequently has little association with rate of ageing, sometimes even lengthening lifespan, has created the impression that ageing has a variety of causes likely too complicated to understand. For fear of getting stuck in this quagmire, the field has moved on to attempting to cure ageing by targeting its observed effects rather than causes, and this has not been without successes. The development of statins to treat atherosclerosis is one example, but as with any treatment aimed at effects rather than causes, such drugs can at best slow the progression of disease, without rejuvenation. For gerontology to progress, we require models of ageing that offer evolutionarily plausible initial causes which can be connected mechanistically to ageing phenotypes and diseases. SDT and metabolic slowdown offer just such a model, including testable predictions and mechanisms which will allow us to assess evidence in the light of competing theories without forcing all data into the tableau of accumulating damage. It is likely that ageing will reflect some combination of selective destruction, other changes from selection (such as pro-growth), and random damage accumulation, which are not exclusive. We suggest that the FEM forms the basis of a model to find the complimentary and contradictory mechanisms of each theory.

## Supporting information

Supplementary model methods, results, and discussion

## Abbreviations Table

| Abbreviation | Full term | Abbreviation | Full term |
| --- | --- | --- | --- |
| **ACC1** | Acetyl-CoA carboxylase 1 | **IR** | Insulin resistance |
| **ACC2** | Acetyl-CoA carboxylase 2 | **k<sub>CM</sub>** | Intrinsic capacity for cellular metabolism |
| **ACoA** | Acetyl-coenzyme A | **L<sub>B</sub>** | Blood lipid |
| **AHCM** | ATP hydrolysing cellular metabolism | **LD** | Lipid droplet |
| **AL** | Ad libitum | **LDL** | Low density lipoprotein |
| **AMP** | Adenosine monophosphate | **LDL-R** | LDL-receptor |
| **AMPK** | AMP-activated protein kinase | **L<sub>i</sub>** | Intracellular lipid |
| **APMM** | ATP producing mitochondrial metabolism | **M** | Mitochondria |
| **ATP** | Adenosine triphosphate | **MAPK** | Mitogen-activated protein kinase |
| **au** | Arbitrary units | **MD** | Mitochondrial dysfunction |
| **βM** | β-cell mass | **mTOR** | Mammalian target of rapamycin |
| **BMI** | Body mass index | **mTORC1** | mTOR complex 1 |
| **CD36** | Cluster of differentiation 36 | **OXPHOS** | Oxidative phosphorylation |
| **CLS** | Crown-like structure | **PGC1α** | Peroxisome proliferator-activated receptor-γ coactivator-1α |
| **CR** | Calorie restriction | **RCT** | Reverse cholesterol transport |
| **DR** | Dietary restriction | **ROS** | Reactive oxygen species |
| **DST** | Disposable soma theory | **SENS** | Strategies for Engineered Negligible Senescence |
| **ECM** | Extracellular matrix | **SDT** | Selective destruction theory |
| **FEM** | Fuel and Energy Model | **s<sub>i</sub>** | Insulin sensitivity |
| **FFA** | Free fatty acid | **TFEB** | Transcription factor EB |
| **GH** | Growth hormone | **TLR4** | Toll-like receptor 4 |
| **IGF-1** | Insulin-like growth factor-1 | **TNF-α** | Tumour necrosis factor-α |
| **IMC** | Intrinsic mitochondrial capacity | **UPR** | Unfolded protein response |

## Author contributions

James Wordsworth developed the theory and model, and wrote the manuscript with input and supervision from Daryl Shanley. Pernille Yde Nielsen helped with coding, modelling, and manuscript development, as well as contributing ideas. Edward Fielder helped develop the falsifiability criteria and along with Sharmilla Chandrasegaran helped with research and contributed to manuscript development.

## Acknowledgements

We would like to thank Professor Tom Kirkwood and Dr Viktor Korolchuk for guidance and input. Images were created using BioRender.com. This work was supported by the Novo Nordisk Fonden Challenge Programme: Harnessing the Power of Big Data to Address the Societal Challenge of Aging (grant number NNF17OC0027812).

