## Supplementary model methods, results, and discussion for "Metabolic slowdown as the proximal cause of ageing and death"

#### STAR Methods

##### Model construction

The Fat & Energy Model (FEM) was constructed using ordinary differential equations (ODEs) based on a non-systematic review of energy homeostasis literature. As with all models, this involved simplifying the biological detail to include only the central effectors of the described processes. This supplementary description of model construction is organised to mirror the layout of the main article, starting with energy dynamics and ending with lipid dynamics.

###### ATP dynamics

***Adenosine triphosphate (ATP) dynamics*** are defined by the processes in equation 1 (shown here to aid interpretation) and described in equation S1.

$$\frac{dATP}{dt} = Glycolysis + APMM - AHCM \quad (1)$$

$$\frac{d[ATP]}{dt} = Glyc + \alpha_3 \cdot IMC \cdot ACoA \cdot ADP \cdot K - \frac{E_{CM} \cdot k_{CM} \cdot \eta(ATP)}{K} \quad (S1)$$

**Glycolysis** is modelled as the parameter **Glyc**, as in equation S1.1, defining the rate at which ATP is produced by the glycolysis of glucose.

$$Glycolysis = Glyc \ (S1.1)$$

Glycolysis is the production of pyruvate from glucose (and here includes the subsequent step producing acetyl-coenzyme A, ACoA, from pyruvate), which also produces a small amount of ATP. We kept Glyc at zero for model simulations, choosing to focus entirely on lipid dynamics in the initial model, so the same parameter could be used for ACoA production (below).

**ATP hydrolysing cellular metabolism** (**AHCM**) was defined as the non-mitochondrial and mitochondrial metabolism requiring the hydrolysis of **ATP** (i.e. the metabolism using up the bioavailable energy, rather than producing it), as in equation S1.2.

$$AHCM = \frac{E_{CM} \cdot k_{CM} \cdot \eta(ATP)}{K} \ (S1.2)$$

We use a single parameter, $\mathbf{k_{CM}}$ to define the basal metabolic rate. The $\mathbf{E_{CM}}$ parameter defines the tissue's upregulation of AHCM in response to external cues (the capacity limited by $k_{CM}$, NB: using two separate parameters allowed us to manipulate these aspects in combination). Within this capacity, AHCM can still be regulated by inhibition via AMP-activated protein kinase (AMPK) activity (K, described below), as is well documented[1,2].

To some extent, AHCM must be dependent on the level of its substrate, ATP. We describe this dependence as a switch function **η(ATP)**, as in equation S1.3. This allows AHCM to occur at maximal (normal) rate while ATP levels are higher than the (low) threshold determined by the parameter, $k_p$, dropping quickly to zero when ATP drops below it.

$$\eta(ATP) = \frac{ATP^{10}}{({k_p}^{10} + ATP^{10})} \qquad (S1.3)$$

**AMP-activated protein kinase** (**AMPK**) **activity** is described by variable **K**, inhibiting AHCM. K is determined by the ratio of **adenosine monophosphate** (**AMP**):ATP, regulated firstly by upstream kinases and phosphatases phosphorylating/dephosphorylating AMPK's activation loop, and secondly by phosphorylation-independent, allosteric kinase activation[3]. While ATP and AMP affect both processes, ADP plays a more minor role, mainly protecting against dephosphorylation without contributing to allosteric activation[4-7]. Therefore, we have only considered AMP and ATP in the activation of AMPK. As shown in equation S1.4, AMPK activation is determined by the ratio of AMP:ATP, while the rate of deactivation increases linearly with active AMPK (corresponding to a spontaneous deactivation process with a fixed average rate).

$$\frac{dK}{dt} = k_{ka}\frac{AMP}{ATP} - k_{kd} \cdot K \quad (S1.4)$$

ATP regulation appears to be highly robust; as described by Hancock, et al. (2006)[8], "the rate of ATP hydrolysis is matched by the rate of ATP supply without measurable changes in ATP over a wide range of ATP demands (up to very intense short-term near-maximal exercise)"[9]. Outside the conditions of intense exercise, AMPK can regulate metabolism and respiration to maintain the correct ratio of ATP:ADP/AMP without affecting the size of the adenosine phosphate (AP) pool.

Adenylate kinase reversibly converts 2 ADP to one AMP and one ATP, maintaining equilibrium levels of ADP and AMP under most conditions[10], which for simplicity we have assumed is a 1:1 ratio so AMP ≈ ADP. This allows us to ignore the least important regulator of K, ADP, as shown in equation S1.5-1.6, where **$ATP_{max}$** is the size of the AP pool, and thus the maximum amount of ATP that could be produced. Thus, the AMP:ATP ratio in equation S1.4 is defined as in equation S1.7.

$$ADP = AMP = \frac{ATP_{max} - ATP}{2} \quad (S1.5)$$

$$ATP = ATP_{max} - 2 \cdot AMP \quad (S1.6)$$

$$\frac{AMP}{ATP} = \frac{ATP_{max} - ATP}{2 \cdot ATP} \quad (S1.7)$$

**ATP producing mitochondrial metabolism** (**APMM**) reflects all metabolism that results in net production of ATP within the mitochondria. Here, we have separated this from the processes of glycolysis and β-oxidation. Glycolysis is usually defined by the production of pyruvate (and 2 ATP); however, for simplification of ACoA dynamics (see equation 2), we have defined both processes to include the following step where pyruvate (and acyl-coenzyme A for β-oxidation) are converted to ACoA, which occurs within the mitochondria. APMM thus refers to utilisation of ACoA in Krebs cycle and the subsequent ATP production during oxidative phosphorylation (OXPHOS) from the NADH and $FADH_2$ it produces.

The factors governing APMM are complex. For example, Krebs cycle can be regulated at multiple points to generate the various intermediates as required for specific metabolic needs other than ATP production. OXPHOS is dependent on the availability of oxygen to accept the electrons at the end of the electron transport chain, forming water by binding the hydrogen ions. For simplicity, we have assumed that external conditions such as the need for Krebs intermediates are constant, and oxygen is in plentiful supply. APMM is thus influenced by the availability of its substrates **adenosine diphosphate** (**ADP**, described by equation S1.5) and ACoA, multiplied by the intrinsic mitochondrial capacity (IMC) to convert ACoA to ATP, and the AMPK activity (K) which stimulates APMM as in equation S1.8.

$$APMM = \alpha_3 \cdot IMC \cdot ACoA \cdot ADP \cdot K \; (S1.8)$$

**Intrinsic mitochondrial capacity** (**IMC**) is modelled as a dynamic variable defined by the functionality of the mitochondrial space per unit volume multiplied by the amount of mitochondrial space (i.e. mass · functionality). In the main article we mention the complex list of effectors influencing the IMC, but in the FEM we focus on K as the primary regulator of all subsequent factors influencing IMC, which therefore determines the rates of Krebs and OXPHOS. Thus, low ATP and high K accelerate Krebs and OXPHOS, and high ATP low K slow them[11,12], and the factors affecting IMC can be condensed into a function of K so that changes in IMC can be defined as in equation S1.9.

$$\frac{d[IMC]}{dt} = IMC\left(\frac{\mu_3}{1+(\frac{\lambda_3}{K})^{\varepsilon_3}} - \frac{\mu_4}{1+(\frac{K}{\lambda_4})^{\varepsilon_4}}\right) \quad (S1.9)$$

The equation is designed to have dynamic compensation with respect to K, forcing K to an equilibrium value determined by the parameters **$\lambda_3$** and **$\lambda_4$** as described by Karin, et al. (2016)[13], discussed below. Thus, the ratio of ATP:AMP and total ATP level are maintained at equilibrium despite changes in metabolism (described further in model discussion).

#### ACoA dynamics

***ACoA dynamics*** are defined by the processes in equation 2 (shown here to aid interpretation) and described in equation S2.

$$\frac{dACoA}{dt} = \beta\text{-}oxidation + Glycolysis - APMM - FFA\ synthesis \quad (2)$$

$$\frac{dACoA}{dt} = \frac{\alpha_2 \cdot \eta_2(L_I) \cdot K \cdot IMC}{\alpha_2 + ACC2 \cdot ACoA^{n_4}} + Glyc - \alpha_3 \cdot IMC \cdot ACoA \cdot ADP \cdot K - ACC1 \cdot ACoA^{n_5} \quad (S2)$$

In addition to glycolysis discussed above, ACoA is produced by β-oxidation, and can either be de-acetylated in APMM or converted back into free fatty acids (FFAs) via FFA synthesis.

ACoA is controlled by robust negative feedback. Excess mitochondrial ACoA is converted to malonyl-CoA by acetyl-CoA carboxylase 2 (ACC2), which then allosterically inhibits the transfer of cytoplasmic acyl-CoA through the mitochondrial membrane into the matrix through carnitine palmitoyltransferase 1 (CPT1, forming acyl-carnitine as an intermediate)[14]. This is frequently the rate-limiting step in **β-oxidation**; however, it is tissue dependent[15]. Thus, the main inhibitory factor for β-oxidation was mitochondrial malonyl-CoA, which we define in equation S2.1 as a function of ACoA.

$$\beta\text{-}oxidation = \frac{\alpha_2}{\alpha_2 + ACC2 \cdot ACoA^{n_4}} \cdot \eta_2(L_I) \cdot K \cdot IMC \quad (S2.1)$$

$$off - switch\ by\ ACoA\text{:} \quad \frac{\alpha_2}{\alpha_2 + ACC2 \cdot ACoA^{n_4}}$$

$$on - switch\ by\ L_I\ (see\ below)\text{:} \quad \eta_2(L_I)$$

$$linear\ for\ K\ and\ IMC$$

K influences the rate of β-oxidation indirectly by activating peroxisome proliferator activated receptor α (PPARα) and peroxisome proliferator-activated receptor-gamma coactivator 1 (PGC1)[16], regulating IMC (equation S1.9), but also directly by inhibiting ACCs[17], and stimulating malonyl CoA decarboxylase (MCD) which degrades malonyl-CoA[18].

It is also dependent on the availability of $L_I$. Just as we described the dependence of AHCM on ATP levels by the function η(ATP), equation S2.2 describes a similar function **$\eta_2(L_I)$** for the dependence of β-oxidation on $L_I$. As most $L_I$ is stored as lipid droplets (LDs), while FFAs are kept relatively constant, the rate of β-oxidation should only become dependent on $L_I$ once LDs have been exhausted and can no longer replenish the FFAs required for APMM.

$$\eta_2(L_I) = \frac{{L_I}^{50}}{({k_l}^{50} + {L_I}^{50})} \qquad (S2.2)$$

Excess mitochondrial ACoA is also converted to citrate (in Krebs cycle) which is then pumped out of mitochondria by tricarboxylate carriers and converted back to ACoA by citrate lyase. Cytoplasmic ACoA then forms the substrate for **FFA synthesis**, converted to malonyl-CoA by acetyl-CoA carboxylase 1 (ACC1, the cytoplasmic version of ACC2). Malonyl-CoA is then converted to FFAs by fatty acid synthase (FAS). Thus, we defined FFA synthesis by equation S2.3, where ACC·ACoA is used as a proxy for malonyl-CoA as for β-oxidation, and constants for ACC1 and FAS are combined to form a single constant (ACC1).

$$FFA\ synthesis = \ ACC1 \cdot ACoA^{n_5} \qquad (S2.3)$$

We have assumed FFA synthesis under simulated conditions is low, setting ACC1 to a value of 0.00001 (compared to a value of 2 for ACC2), reflecting that we are initially modelling lipid dynamics in the absence of alternative fuels which would be converted to fats, so FFA synthesis will be downregulated.

#### Lipid dynamics

***Lipid dynamics*** have been simplified to consider only FFAs, triglycerides (three fatty acids bonded to glycerol), and the macromolecules and organelles which bind and carry them. FFAs in **meals** enter the body via the gut, are esterified to form triglycerides, and transferred to the blood attached to lipoproteins (chylomicrons from gut). Equally, FFAs from internal sources are esterified and bound to very-low-density lipoprotein (VLDL) in the liver, which is transported around the blood forming low-density lipoprotein (LDL) and high-density lipoprotein (HDL). Unlike glucose, FFAs are not soluble in water, so only a tiny percentage (<0.01%) are carried free in the plasma with >95% bound to albumin[19], while more than 90% of total fatty acids in plasma are stored as esters (such as triglycerides), which are mainly transported on lipoproteins. This combined pool of **blood lipids** is referred to as **$L_B$**. Similarly, the combined pool of FFAs, triglycerides, and lipid droplets (LDs) are grouped together as **intracellular lipid** (**$L_I$**).

##### *$L_B$ dynamics*

Changes in $L_B$ are defined by the processes in equation 3 (shown here to aid interpretation) and described in equation S3.

$$\frac{dL_B}{dt} = Meal - Import + Export \quad (3)$$

$$\frac{dL_B}{dt} = Meal - L_B(C_1 + I \cdot s_i) - \frac{\eta_2(L_I)}{(C_2 + \ s_i * I)^{n_1}} \qquad (S3)$$

The **import** of $L_B$ is controlled primarily by **insulin** (variable **I**) through the activation of cluster of differentiation 36 (CD36), in a similar mechanism to glucose transport via glucose transporter type 4

(GLUT4)[20-22]. Insulin also stimulates lipoprotein lipase in the wall of the vascular endothelium, which breaks the ester bonds of triglycerides to release the FFAs for import to adipose tissue[23]. The degree to which insulin stimulates the import of $L_B$ is dependent firstly on $L_B$ and secondly on the insulin sensitivity, $s_i$ (variable described below) of the adipocytes. Thus, import is described as in equation S3.1, with insulin-independent import defined by the parameter **$C_1$**.

$$Import = L_B(C_1 + I \cdot s_i) \quad (S3.1)$$

Due to the similarities between $L_B$ and blood glucose regulation, we modelled import of $L_B$ with a derivation of the blood glucose model produced by Topp, et al. (2000)[24], called the β-cell, insulin, glucose (BIG) model. We adapted a derivation of the BIG model produced by Karin, et al. (2016)[13] for import and insulin dynamics, substituting $L_B$ for glucose as in equation S3.2, reflecting that insulin is produced by β-cells in the pancreas, which are stimulated by circulating FFAs both to produce insulin[25,26] and proliferate[27-29] similarly to blood glucose. Thus, as in equation S3.2, insulin is produced in an $L_B$-dependent manner at rate **q** per β-cell, multiplied by the number of β-cells, or **β-cell mass** (variable **βM**). Insulin then degrades spontaneously at rate **$\gamma_i$**.

$$\frac{dI}{dt} = q \cdot \beta M \cdot {L_B}^2 - \gamma_i \cdot I \quad (S3.2)$$

This model was appealing because Karin, et al. (2016)[13] demonstrated that the BIG model had dynamic compensation, meaning that for any value of q or $s_i$ the system would adapt to return $L_B$ to equilibrium (see model discussion for further detail). This occurs because of the βM expanding and contracting according to the level of $L_B$ as in equations S3.3-3.4, based on those used by Karin, et al. (2016)[13]. In the FEM, $L_B$ stimulates proliferation and inhibits apoptosis so that βM grows and shrinks according to insulin production requirements. It forces $L_B$ to an equilibrium value determined by the parameters **$\lambda_1$** and **$\lambda_2$**.

$$\frac{d[\beta M]}{dt} = \beta M(Proliferation\ rate - Apoptosis\ rate) \quad (S3.3)$$

$$\frac{d[\beta M]}{dt} = \beta M\left(\frac{\mu_1}{1 + (\frac{\lambda_1}{LB})^{\varepsilon_1}} - \frac{\mu_2}{1 + (\frac{LB}{\lambda_2})^{\varepsilon_2}}\right) \quad (S3.4)$$

The BIG model did not consider $L_I$ **export**, but evidence suggests this is also dependent on insulin. Insulin inhibits lipolysis and export, inhibiting lipolysis in an AKT-independent[30] and dependent mechanism by dephosphorylating hormone sensitive lipase and preventing de-esterification of intracellular triglycerides in LDs[31,32]. We modelled export with an insulin-dependent and independent component, **$C_2$**, as described in equation S3.5.

$$Export = \frac{\eta_2(L_I)}{(C_2 + s_i * I)^{n_1}} \quad (S3.5)$$

The $\eta_2(L_I)$ function limiting β-oxidation when $L_I$ drops below the threshold will also apply to export, as the central function of export (from the adipose) is to replenish $L_B$ and ensure the supply of FFAs for *non-adipose* β-oxidation (as well as broader metabolic needs not considered here). Notably, as most non-adipose tissues cannot store $L_I$ as LDs, excess $L_I$ is toxic[33-35], necessitating swift return to equilibrium. The $\eta_2(L_I)$ function may thus not apply to non-adipose export, while import likely adapts much more swiftly with alterations to $s_i$. As the functions for non-adipose export and import likely differ from the functions for adipose tissue, $L_B$ would more accurately be described by equation S3.6.

$$\frac{dL_B}{dt} = Gut\ absorption - Adipose\ Import + Adipose\ Export - Non\text{-}adipose\ import + Non\text{-}adipose\ export \quad (S3.6)$$

As there is little storage of $L_I$ in non-adipose tissues, the level of $L_I$ import will roughly equal the level of β-oxidation over time, either because excess $L_I$ is exported back into the blood or the insulin sensitivity of the tissues quickly adapts to reduce import to match their metabolic rate. Therefore, net import ≈ β-oxidation. Thus, equation 3.6 could be re-written as in equation S3.7, and **Meal** defined as in equation S3.8.

$$\frac{dL_B}{dt} = Gut\ absorption - Adipose\ Import + Adipose\ Export - Non\text{-}\ adipose\ \beta\text{-}\ oxidation \quad (S3.7)$$

$$Meal = Gut\ absorption - Non\text{-}\ adipose\ \beta\text{-}\ oxidation \quad (S3.8)$$

The implications of this simplification are described in the model discussion, but it allowed us to limit the FEM to the adipose and blood rather than including additional tissues. NB: we used the term 'Meal' in the main text as this was sufficiently descriptive for the textual analysis of metabolic slowdown before considering that the FEM separates adipose from non-adipose dynamics.

Thus, when we refer to the **insulin sensitivity** (variable $\mathbf{s_i}$) we refer exclusively to the adipose. Although an in-depth review of $s_i$ regulation is beyond the scope of the FEM, it is worth noting that because adipose can store $L_I$ (as LDs) it cannot regulate $s_i$ by the size of the FFA pool as can other tissues. To avoid bursting, adipocytes must regulate $s_i$ with respect to their $L_I$ as stored in LDs. It is well established that small adipocytes are more insulin sensitive than large ones[36,37], and adipocyte size correlates with $s_i$ independent of fat mass[38].

One mechanism reducing $s_i$ results from the redistribution of cholesterol from the plasma membrane to the LDs[39,40]. Cholesterol forms a major part of lipid rafts called caveolae which act as major docking sites for insulin sensitive transporters such as CD36 and GLUT4[41,42]. Therefore, cholesterol depletion of the membrane prevents endocytosis and insulin signalling in adipocytes[43], promoting insulin resistance (IR). Importantly, the main protein stabilising caveolae, caveolin 1 (CAV1), regulates expansion of LDs by its transfer from the plasma membrane to the membrane of the organelles[44,45], thus as LDs expand, adipocytes become less insulin sensitive. Therefore, $s_i$ is reduced by increasing $L_I$ as defined by equation S3.9, while production of insulin receptor was modelled to occur at a constant rate defined by parameter, $\mathbf{k_s}$.

$$\frac{ds_i}{dt} = k_s - \frac{s_i \cdot k_r \cdot {L_I}^{n_2}}{1 + {L_I}^{n_2}} \quad (S3.9)$$

**Plaque formation** and growth was modelled as a function of elevated $L_B$. We suggested there would be an **equilibrium point** where the rate of plaque degradation equalled the rate of plaque production, $\mathbf{P_{eq}}$, and risk from plaques would change as shown in equation S3.10 (NB: risk was reset to 1 at time t = 1900 (after model has reached equilibrium), and could not drop below zero (see supplementary code, lines 61-72).

$$\frac{d[Risk]}{dt} = R_R(L_B - P_{eq}) \quad (S3.10)$$

When $L_B$ rises above $P_{eq}$, $(L_B - P_{eq} > 0)$ risk increases as the plaques enlarge and spread (linearly as the distance from equilibrium increases), and when $L_B$ drops back below $P_{eq}$ the plaques decrease in size and number as the removal mechanisms begin to work faster than lipid deposition. Notably, $P_{eq}$ is not (necessarily) the same as the equilibrium value for $L_B$, and does not have to be constant. We could incorporate 'wear and tear' induced endothelial dysfunction as a steadily declining $P_{eq}$, or $P_{eq}$ could be a function of inflammation and/or insulin resistance; see Theofilis, et al. (2021)[46] for recent review.

##### $L_I$ dynamics

Changes in $L_I$ are defined by the processes in equation 4 (shown here to aid interpretation) and described in equation S4.

$$\frac{dL_I}{dt} = Import - Export + FFA\ synthesis\ - \beta\text{-}\ oxidation - Apoptosis\ \ (4)$$

$$\frac{dL_I}{dt} = L_B(C_2 + I \cdot s_i) - \frac{\eta_2(L_I)}{(C_2 +\ s_i * I)^{n_1}} + ACC1 \cdot ACoA^{n_5} - \frac{\alpha_2 \cdot \eta_2(L_I) \cdot K \cdot IMC}{\alpha_2 + ACC2 \cdot ACoA^{n_4}} - \frac{k_a \cdot L_I \cdot T}{(C_3 +\ s_i \cdot I)^{n_3}} \quad (S4)$$

The processes for Import, Export, FFA synthesis and beta-oxidation have all been described above.

**Apoptosis** removes hypertrophic cells via activation of a localised immune response and inflammation. Hypertrophic cells produce inflammatory cytokines which attract immune cells such as macrophages[47] and cause a phenotypic switch toward the inflammatory M1 polarisation[48].

Basal inflammation in the adipose tissue is a highly complex process, including an array of activatory and inhibitory cytokines and chemokines, as well as T and B cells in addition to macrophages and monocytes, some of which can also function to inhibit inflammation as well switch between pro and anti-inflammatory states. For the FEM, we modelled inflammation by simplifying it to a single variable, **TNF-α** (**T**), which is thought to be responsible for the extrinsic apoptosis of hypertrophic adipocytes[49,50] (see model discussion for implications). Hence knockout mouse models with increased inflammation and apoptosis have reduced fat mass[51].

Infiltrating macrophages localize around large or dead adipocytes forming crown-like structures[52], suggesting that hypertrophic adipocytes are preferentially targeted for apoptosis, perhaps in part because they are insulin resistant[50,53,54]. Also in obese children, the size of fat cells correlated more strongly with inflammatory markers than either body mass index (BMI) or fat mass[38]. Therefore, we modelled apoptosis as dependent on $L_I$, representing the number of hypertrophic cells, and degree of inflammation, T, as in equation S4.1.

$$Apoptosis = \frac{k_a * L_I * T}{(C_3 +\ s_i * I)^{n_3}} \quad (S4.1)$$

Hypertrophic cells have increased expression of Toll-like receptor 4 (TLR4) and NF-κB expression which not only attract macrophages[55], but sensitize cells to apoptosis[56,57], and contribute to IR[58,59]. Importantly, death-receptor-induced apoptosis is inhibited by IGF-1 and insulin in white[50,60] and brown[61] adipocytes. Insulin promotes survival in multiple cell types[53,54], and increasing insulin concentrations in vitro protect adipocytes from TNF-α induced apoptosis[50]. Thus, while both hypertrophic adipocytes and infiltrating macrophages cause adipose inflammation, mainly the insulin resistant hypertrophic adipocytes undergo apoptosis[36-38] (Figure 2D, blue). Thus, apoptosis was inhibited by insulin dependent on the degree of $s_i$ (and independently of insulin, **$C_3$**).

We modelled variable **T** in equation S4.2 to be increased primarily by the number of hypertrophic cells producing inflammatory T, represented by $L_I$. The rate of T degradation was modelled as spontaneous, similarly to insulin degradation, dependent on the amount of T multiplied by degradation constant, **$\gamma_t$**.

$$\frac{dT}{dt} = \frac{{L_I}^H}{{\alpha_1}^H +\ {L_I}^H} - \gamma_t \cdot T\ (S4.2)$$

#### Model running and storage

The FEM was constructed using R version 3.6.3 with the deSolve package for ordinary differential equations (ODEs). The R code for the model and all data analysis are available on FAIRDOM Hub with the following identifying links:

DOI: 10.15490/fairdomhub.1.model.851.1 SEEK ID: https://fairdomhub.org/models/851?version=1

Values and definitions for static parameters are found in Table S1, and initial variable values and adjusted parameters in Table S2. However, it should be noted that the FEM is not supposed to represent any specific biological conditions but rather broad biological contexts. It is designed to show qualitative outcomes which determine whether systems are in homeostasis and the directional changes of variables relative to others. The parameter values and any simulated quantities are therefore mainly arbitrary, as are the units (au).

| Parameter | Definition | Value |
|---|---|---|
| Lipid dynamics | | |
| C1 | Insulin independent import of $L_B$ | 0.001 |
| C2 | Insulin independent import of $L_I$ | 0.001 |
| C3 | Insulin independent inhibition of apoptosis | 0.001 |
| $n_1$ (n1) | Exponent of insulin-dependent and independent inhibition of export | 2 |
| $n_3$ (n3) | Exponent of insulin-dependent and independent inhibition of apoptosis | 1 |
| $n_4$ (n4) | Exponent of formation of malonyl-CoA by ACC2, inhibiting β-oxidation | 5 |
| $n_5$ (n5) | Exponent of formation of malonyl-CoA by ACC1, stimulating FFA synthesis | 5 |
| $\alpha_2$ (a2) | Constant for switch function of ACoA on β-oxidation | 625 |
| $k_a$ (ka) | Rate constant for $L_I$ and T induced apoptosis | 0.02 |
| $k_l$ (kl) | Threshold value for $L_I$ in export and β-oxidation | 0.5 |
| ACC1 | Enzyme catalysing conversion of cytoplasmic ACoA into malonyl-CoA, stimulating FFA synthesis | 0.00001 |
| ACC2 | Enzyme catalysing conversion of mitochondrial ACoA into malonyl-CoA, inhibiting β-oxidation | 2 |
| $P_{eq}$ (Eq) | Equilibrium where plaque formation equals removal | 1.15762 |
| $R_R$ (Rr) | Rate of change in plaque-associated risk | 1 |
| Insulin dynamics | | |
| q | Rate constant for insulin production | 0.01 |
| $\gamma_1$ (gami) | Degradation rate of insulin | 0.05 |
| $\mu_1$ (mu1) | Rate constant for βM increase (proliferation) | 0.021 |
| $\mu_2$ (mu2) | Rate constant for βM decrease (apoptosis) | 0.025 |
| $\lambda_1$ (lam1) | $L_B$ threshold for increasing βM (proliferation) | 1.556 |
| $\lambda_2$ (lam2) | $L_B$ threshold for decreasing βM (apoptosis) | 0.8 |
| $\varepsilon_1$ (eta1) | Exponent for $L_B$-induced βM proliferation | 10 |
| $\varepsilon_2$ (eta2) | Exponent for $L_B$-inhibition of βM apoptosis | 8.5 |
| $k_s$ (ks) | Rate of increasing insulin sensitivity | 0.5 |
| $k_r$ (kr) | Rate $L_I$ reduces insulin sensitivity | 1 |
| $n_2$ (n2) | Exponent for reduction of insulin sensitivity by $L_I$ | 0.01 |
| Inflammation | | |
| H | Exponent for the induction of inflammation by $L_I$ | 7 |

| | | |
|---|---|---|
| $\alpha_1$ (a1) | Constant for switch function of $L_I$ on inflammation | 100 |
| $\gamma_t$ (gamt) | Degradation rate of TNF-α | 0.5 |
| ATP dynamics | | |
| $\alpha_3$ (a3) | Rate constant for APMM | 0.2 |
| $k_{ka}$ (kka) | Activation rate of AMPK | 1 |
| $k_{kd}$ (kkd) | Deactivation rate of AMPK | 1 |
| $k_p$ (kp) | Threshold value for ATP in AHCM | 1 |
| $ATP_{max}$ | Size of adenosine phosphate pool | 100 |
| $\mu_3$ (mu3) | Rate constant for IMC increase | 0.021 |
| $\mu_4$ (mu4) | Rate constant for IMC decrease | 0.025 |
| $\lambda_3$ (lam3) | AMPK activity threshold for increasing IMC | 10.892 |
| $\lambda_4$ (lam4) | AMPK activity threshold for decreasing IMC | 5.6 |
| $\varepsilon_3$ (eta3) | Exponent for K-induced increase in IMC | 1.7 |
| $\varepsilon_4$ (eta4) | Exponent for K-inhibition of IMC decrease | 8.5 |
| Glyc (G) | Rate of glycolysis | 0 |

*Table S1 | Parameter values for the FEM. Terms in brackets are the parameter names used in the code.*

| Variable | Symbol | Full name | Initial value |
|---|---|---|---|
| ATP | A | ATP | 16.66667 |
| ACoA | Aco | Acetyl-CoA | 5 |
| βM | BM | Β-cell mass | 1 |
| I | I | Insulin | 2.3955 |
| IMC | Ma | Intrinsic mitochondrial capacity | 0.04 |
| $L_B$ | LB | Blood lipid | 5 |
| $L_I$ | LI | Intracellular lipid | 100 |
| K | K | AMPK activity | 5 |
| $s_i$ | si | Insulin sensitivity | 0.9656780 |
| T | TN | TNF-α | 0 |
| **Parameter** | **Symbol** | **Definition** | **Initial Value and changes** |
| $E_{CM}$ | ECM | Elevation of AHCM | 1 → 1.5 simulates exercise |
| $k_{CM}$ | MR | Basal metabolic rate | 1, see equation S6 simulating metabolic slowdown. |
| Meal | m | Meal | 1.2 → 1.201 simulates CR to AL feeding |

*Table S2 | Variables and parameters which change during model simulations. Symbol refers to the names used in the code.*

Parameter estimation involved changing each parameter ±1,2,5, and 10% and plotting the effects of changes over time (data not shown). Then to compare effects of each parameter on the variables and processes, we firstly filtered the data to time ≥2000 au (removing the initial equilibration effects) and then used the data for ±10% change in each parameter. We calculated the difference between 10% increase and decrease for each timepoint and took the mean difference across all timepoints for each variable and process, termed **X**. Then we normalised X values by dividing by the absolute maximum X value for each variable (i.e. the value for the parameter with greatest impact on the variable), as shown in equation S5.

$$norm(X) = \frac{X}{\max(|X|)} \qquad (S5)$$

### Results

#### Parameter estimation at equilibrium

We ran initial simulations of the FEM for 2000 arbitrary units (au) to ensure all variables and processes reached equilibrium. We used the parameter values in Table S1 and starting variable values and $E_{CM} = 1$, $k_{CM} = 1$, and Meal = 1.2 from Table S2.

As all variables and processes had reached equilibrium long before t=1900 au, for all future data we removed t<1900 au so the dynamics could be compared (visually and numerically) to equilibrium values rather than including the equilibration effects. To test whether the model was performing appropriately, we ran a parameter estimation as described in the model running and storage, using a 10% change in parameter value to observe effects on each variable and process. Figure S1 shows the relative impact of changing parameters (±10%) for each variable or process: all values are shown as a fraction of the parameter with the greatest impact (by decreasingly bright colour). Red indicates that increasing the parameter has a positive effect on the variable/process and blue a negative effect.

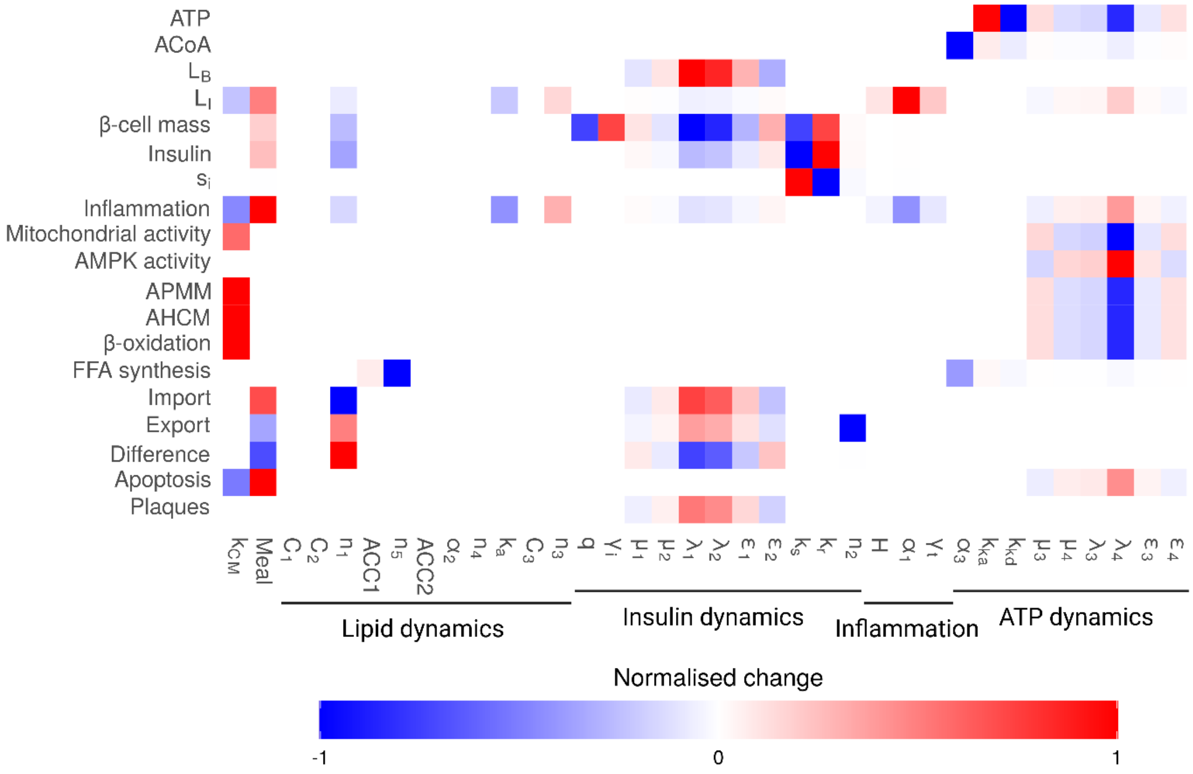


*Figure S1 | Parameter estimation for FEM post equilibration. Values were normalised as described in the model running and storage section so that each value is relative to other parameters for each individual variable/process. An absolute value of 1 (or -1) represents the parameter that has the greatest effect on that variable or process. Positive values reflect that increases in the parameter increase the variable or process, while negative values reflect that increases in the parameter decrease the variable or process. Abbreviations: ACoA, acetyl-CoA; AHCM, ATP hydrolysing metabolism; ATP, adenosine triphosphate; APMM, ATP producing mitochondrial metabolism; $L_B$, blood lipid; $L_I$, intracellular lipid; $s_i$, insulin sensitivity.*

The parameters for ATP dynamics affected most variables and processes, except those which were instead mainly affected by insulin dynamics. Both ATP dynamics and insulin dynamics influenced inflammation (TNF-α) and the parameters for inflammation mainly impacted $L_I$.

Six lipid dynamics parameters affected no variables or processes (at the level of 10% change). These included $C_1$, $C_2$, and $C_3$, which reflected the insulin independent components of import, export (inhibition), and apoptosis (inhibition), respectively. This is an interesting observation as these processes can be insulin-independent[62,63], and the $C_1$ parameter for insulin independent import was adopted from the BIG model equation for glucose dynamics used by Karin, et al. (2016)[13]. We predict as the FEM is expanded with additional variables and processes, these parameters will gain relevance. Additionally, the parameters $\alpha_2$, ACC2, and $n_4$ also had no effect on any variables or processes. All these parameters are part of the β-oxidation equation (S2.1), but have no impact on the FEM because they do not affect the rate of β-oxidation, which is instead determined by the rate of AHCM through $k_{CM}$ and the parameters which affect the equilibrium of ATP ($\mu_3$, $\mu_4$, $\lambda_2$, $\lambda_4$, $\varepsilon_3$, and $\varepsilon_4$), as shown in Figure S1. We therefore tested if these parameters would change under conditions of metabolic slowdown.

#### Model behaviour to stepwise changes in metabolic rate and meal

First, we made sure the model responded appropriately to single step changes in $k_{CM}$ (defining the basal metabolic rate and intrinsic capacity for AHCM) and meal values, as shown in Figure S2.

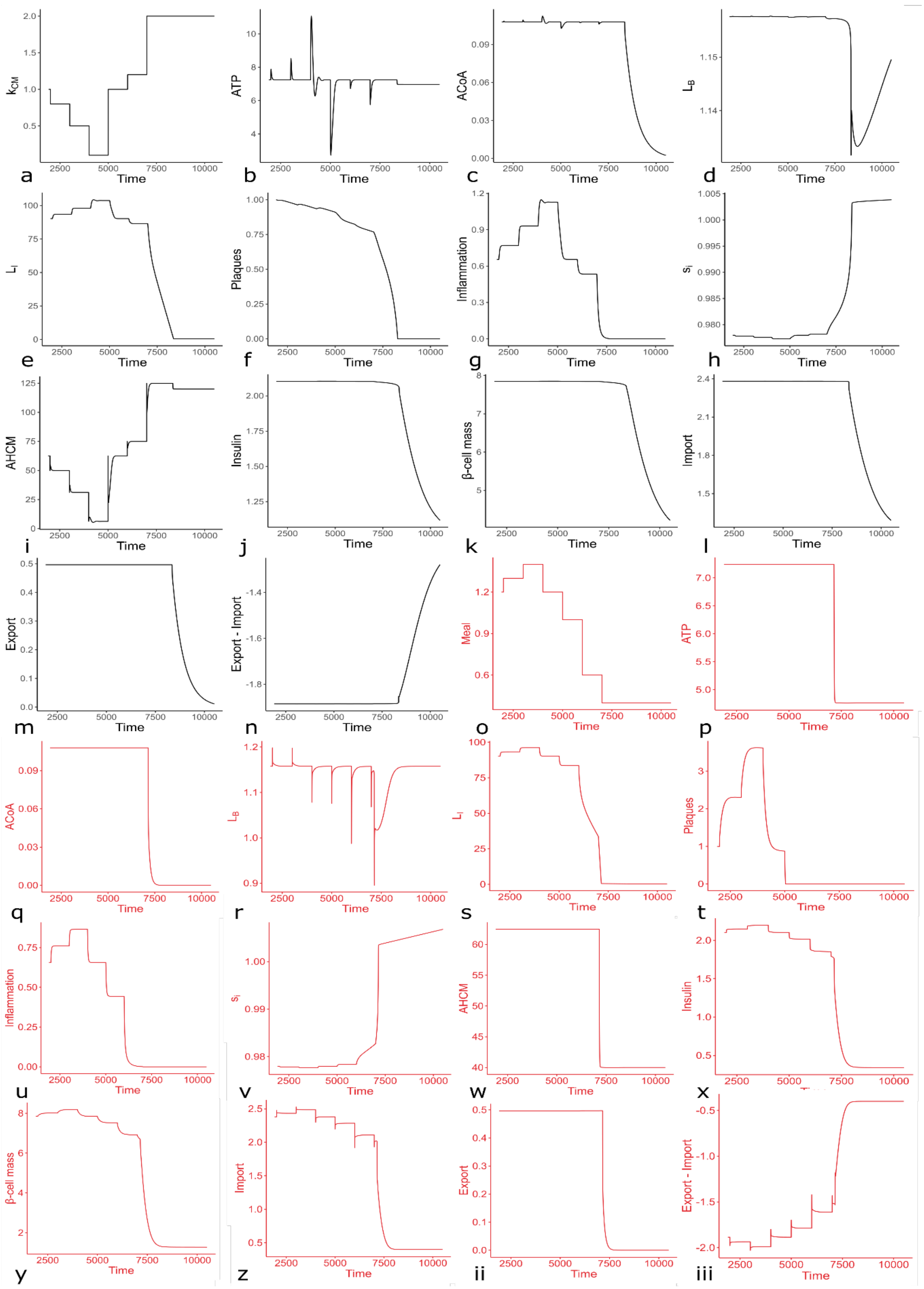

*Figure S2 | Effects of step changes in $k_{CM}$ (black) and Meal (red) on key parameters. (A-N) Effects of step changes in $k_{CM}$. (A) Input changes in $k_{CM}$. (B) ATP. (C) ACoA. (D) $L_B$. (E) $L_I$. (F) Plaques. (G) Inflammation. (H) $s_i$. (I) AHCM. (J) Insulin. (K) β-cell mass. (L) Import. (M) Export. (N) Export minus import. (O-iii) Effects of step changes in Meal. (O) Input changes in Meal. (P) ATP. (Q) ACoA. (R) $L_B$. (S) $L_I$. (T) Plaques. (U) Inflammation. (V) $s_i$. (W) AHCM. (X) Insulin. (Y) β-cell mass. (Z) Import. (ii)*

*Export. (iii) Export minus import. Abbreviations: ACoA, acetyl-CoA; AHCM, ATP hydrolysing metabolism; ATP, adenosine triphosphate; $L_B$, blood lipid; $L_I$, intracellular lipid; $s_i$, insulin sensitivity.*

The results indicated that single changes in $k_{CM}$ and Meal caused appropriate changes in all variables and processes as suggested by the literature and described in the model construction. ATP, ACoA, and $L_B$ all retained a single equilibrium value despite changes in metabolic rate and meal, suggesting the FEM could robustly maintain these equilibria in response to changes in diet and metabolism. Equilibrium $L_I$ shifted as predicted from the data suggesting stored $L_I$ (and weight) has no fixed value per individual and shifts according to metabolic and dietary factors[64].

Importantly, the equilibria for ATP, ACoA, and $L_B$ only shifted when either metabolic rate (and thus $L_I$ burning) increased above the capacity of the (fixed) Meal value to replenish it, or the Meal value decreased below the necessary (fixed) rate of AHCM to maintain cellular function. Both caused the depletion of $L_I$ below the threshold determined by equation S2.2, at which point the model had no remaining stored fat to replenish $L_B$ or utilise in metabolism. In reality, this would reflect death by starvation. There are notably multiple mechanisms to supply fuel for the AHCM of crucial tissues as $L_I$ (and other readily available fuel) borders exhaustion, which mainly involve the catabolism of less important tissues[65]. As the FEM has not been built to address these pathways, they are not included, and the simulations suggested that all variables and processes responded appropriately at least up to the point of $L_I$ exhaustion. We therefore modelled continuous metabolic slowdown.

#### Model behaviour to continuous gradual AHCM slowdown

Metabolic slowdown was modelled as a continuous decline in $k_{CM}$ over time (from t=2000 au), as defined by equation S6.

$$\frac{dk_{CM}}{dt} = -\frac{k_{CM}}{4000} \quad (S6)$$

The results for key variables and processes are shown in Figure 4 and discussed in the main article. We did an additional parameter estimation for the model during metabolic slowdown and the results were the same as in Figure S1 (data not shown). As was the case for repeated shifts in $k_{CM}$ (Figure S2), the FEM responded appropriately to continually decreasing $k_{CM}$, except that β-cell mass (βM) and $L_I$ showed logarithmic growth (Figure 4F and J).

#### Model behaviour with limited organ growth

In reality both adipose mass (i.e. $L_I$ storage capacity) and βM would be finite, limited by organ size. Notably, the same is true for excreting excess $L_B$ via the liver in the bile (reverse cholesterol transport, RCT). To represent all these finite capacities, we added a maximal βM to the FEM (as it already included βM and not liver or adipose mass) as in equation S7.

$$\frac{d[\beta M]}{dt} = 0 \qquad If\left(\frac{\mu_1}{1 + (\frac{\lambda_1}{LB})^{\varepsilon_1}} > \frac{\mu_2}{1 + (\frac{LB}{\lambda_2})^{\varepsilon_2}} AND\ \beta M \geq Threshold\right) \quad (S7)$$

Once βM reached the threshold (7.849) it could no longer increase, preventing the necessary increase in insulin production to sustain $L_B$ as AHCM continued to decline. To supplement the results shown in Figure 4J-L, we included an additional scenario in simulations where the declining $k_{CM}$ from t = 2000 was reversed at t = 6250 (after the βM plateau) to show that βM could still decrease from maximal size as required for homeostasis. The results for key variables are shown in Figure S3. ACoA mirrors ATP (not shown).

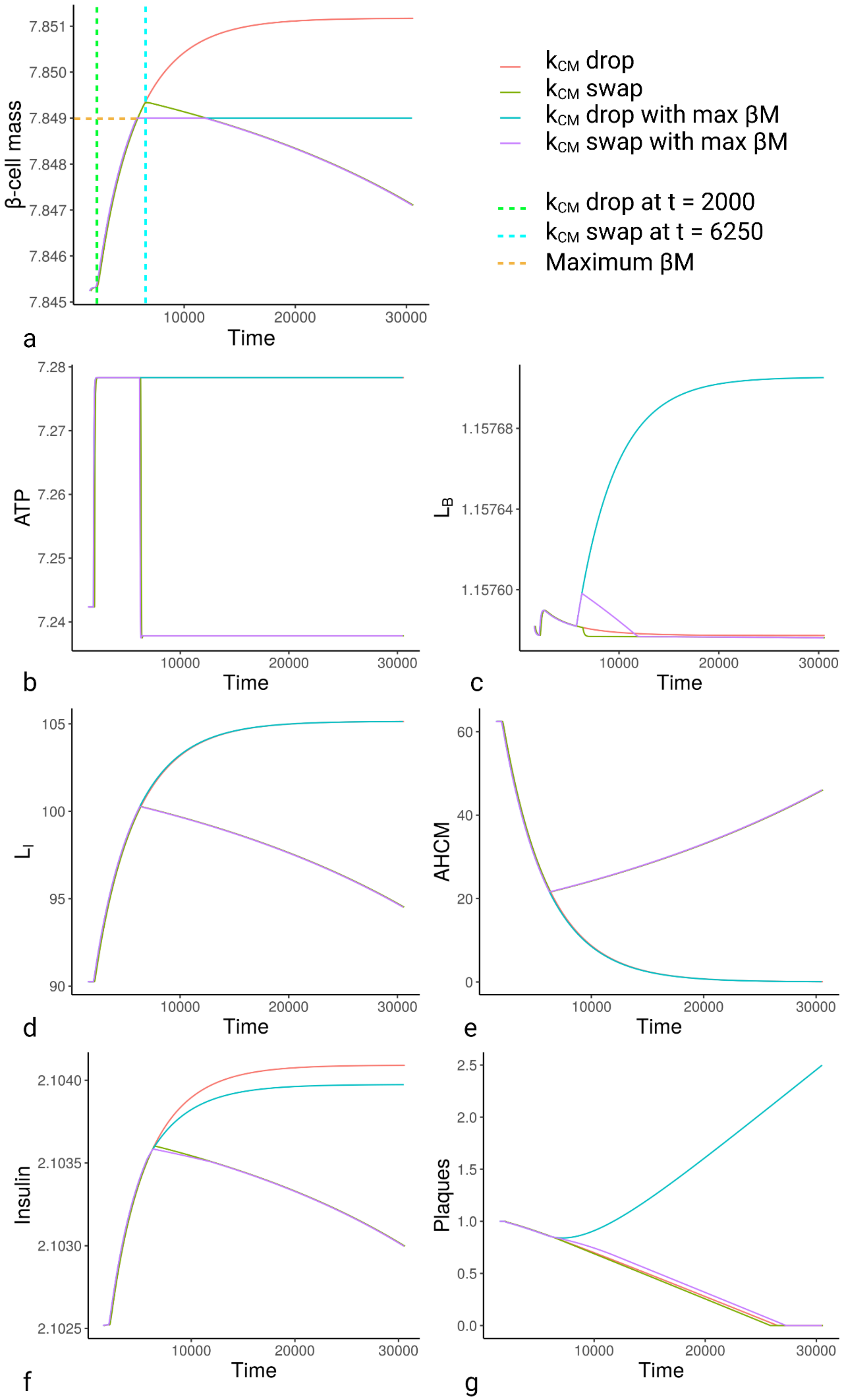

β-cell mass
Time
a
kCM drop
kCM swap
kCM drop with max βM
kCM swap with max βM
kCM drop at t = 2000
kCM swap at t = 6250
Maximum βM
ATP
Time
b
LB
Time
c
LI
Time
d
AHCM
Time
e
Insulin
Time
f
Plaques
Time
g

*Figure S3 | Effect of maximum value of βM on key variables. To demonstrate the effects of a maximum value of βM of 7.849 au, we simulated declining metabolic rate as in Figure 4A ($k_{CM}$ drop) with and without the max βM. To demonstrate that βM could decrease from this mass as required by energy homeostasis, we also simulated a rise in $k_{CM}$ after the maximum βM had been reached ($k_{CM}$ swap). As some simulations were identical, we added +100 au to the time of $k_{CM}$ drop and $k_{CM}$ swap so they could be distinguihsed from the other lines on the graphs.* (A) β-cell mass. (B) ATP. (C) $L_B$. (D) $L_I$. (E) AHCM. (F) Insulin. (G) Plaques. *Abbreviations: AHCM, ATP hydrolysing metabolism; ATP, adenosine triphosphate; $L_B$, blood lipid; $L_I$, intracellular lipid; $s_i$, insulin sensitivity.*

A further parameter estimation of the FEM including a maximum βM demonstrated that the rate of insulin production, q, and the rate of insulin degradation, $\gamma_i$, gained additional significance affecting multiple variables and processes, as did $k_s$ and $k_r$ determining the rate of change in insulin sensitivity, as shown in Figure S4A (compared to Figure S1).

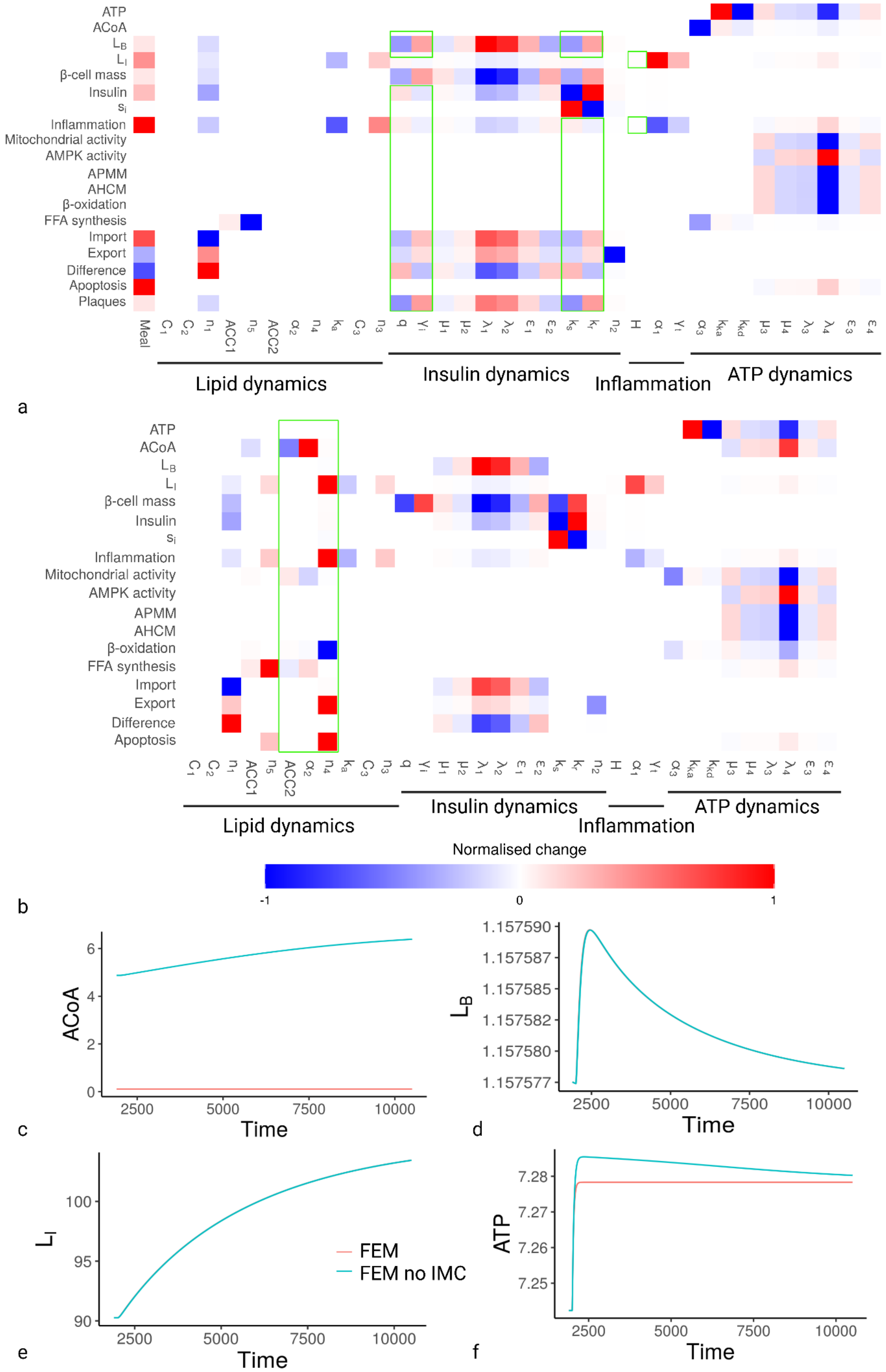

ATP
ACoA
$L_B$
$L_I$
β-cell mass
Insulin
$s_i$
Inflammation
Mitochondrial activity
AMPK activity
APMM
AHCM
β-oxidation
FFA synthesis
Import
Export
Difference
Apoptosis
Plaques
Meal
ACC1
ACC2
Lipid dynamics
Insulin dynamics
Inflammation
ATP dynamics
a
b
Normalised change
-1
0
1
ACoA
Time
c
$L_B$
d
$L_I$
e
FEM
FEM no IMC
ATP
f

*Figure S4 | Parameter estimation and variables for modified FEMs. (A) FEM undergoing metabolic slowdown with maximum βM of 7.849 au. (B) FEM without IMC regulation of β-oxidation. (A-B) Values were normalised as described in the model running and storage so that each value is relative to other parameters for each individual variable/process. An absolute value of 1 (or -1) represents the parameter that has the greatest effect on that variable or process. Positive values reflect that increases in the parameter increase the variable or process, while negative values reflect that increases in the parameter decrease the variable or process. Green boxes show differences to parameter estimation under basal conditions.* (C-F) Simulations of FEM *without IMC regulation of β-oxidation*. (C) ACoA. (D) $L_B$. (E) $L_I$. (F) ATP. *Abbreviations: ACoA, acetyl-CoA; AHCM, ATP hydrolysing metabolism; ATP, adenosine triphosphate; APMM, ATP producing mitochondrial metabolism; FEM, Fuel and energy model; IMC, intrinsic mitochondrial capacity; $L_B$, blood lipid; $L_I$, intracellular lipid; $s_i$, insulin sensitivity.*

#### Model behaviour when β-oxidation is independent of IMC

Notably, a maximum βM did not affect the parameters $\alpha_2$, ACC2, and $n_4$ which were part of the β-oxidation equation (S2.1), which still had no effect on any variables or processes. Importantly, these parameters govern how ACoA impacts β-oxidation, and ACoA levels are in equilibrium during metabolic slowdown. To test whether these parameters were significant to the FEM under conditions where ACoA was not in equilibrium, we changed the β-oxidation equation (S2.1) to exclude the impact of IMC, as shown in equation S8.

$$\beta\text{-}oxidation = \frac{\alpha_2 \cdot \eta_2(L_I) \cdot K \cdot IMC}{\alpha_2 + ACC2 \cdot ACoA^{n_4}} \rightarrow \frac{\alpha_2 \cdot \eta_2(L_I) \cdot K}{\alpha_2 + ACC2 \cdot ACoA^{n_4}} \qquad (S8)$$

As indicated by the results in Figure S4B, all three parameters had multiple impacts on variables and processes when IMC no longer impacted β-oxidation, although only $n_4$ affected the rate of β-oxidation itself, while ACC2 and $\alpha_2$ primarily impacted levels of ACoA, FFA synthesis, and IMC. Removing IMC from the β-oxidation equation demonstrates the robustness of the FEM to maintain the various equilibria: although ACoA levels rise, this does not impact $L_B$, or $L_I$, and ATP still returns to equilibrium, although more slowly (Figure S4C-F).

#### The impact of metformin on ageing as modelled by the FEM

To model the impact of metformin we altered several parameters, firstly, $k_{ka}$ and $k_{kd}$ which determine the rates of AMPK activation and deactivation respectively. We simulated two doses of metformin, introduced between t = 2500 and t = 5000 au and then between t = 7500 and t = 10000 au, either by increasing $k_{ka}$*1.5 or reducing $k_{kd}$/1.5 to represent increased rate of activation and decreased rate of deactivation, respectively. The results were identical for changes in $k_{ka}$ and $k_{kd}$ (latter not shown). Increasing $k_{ka}$ and its effects on $L_B$, plaques, AMPK activity, ATP, inflammation, AHCM, and APMM are shown in Figure S5A-D.

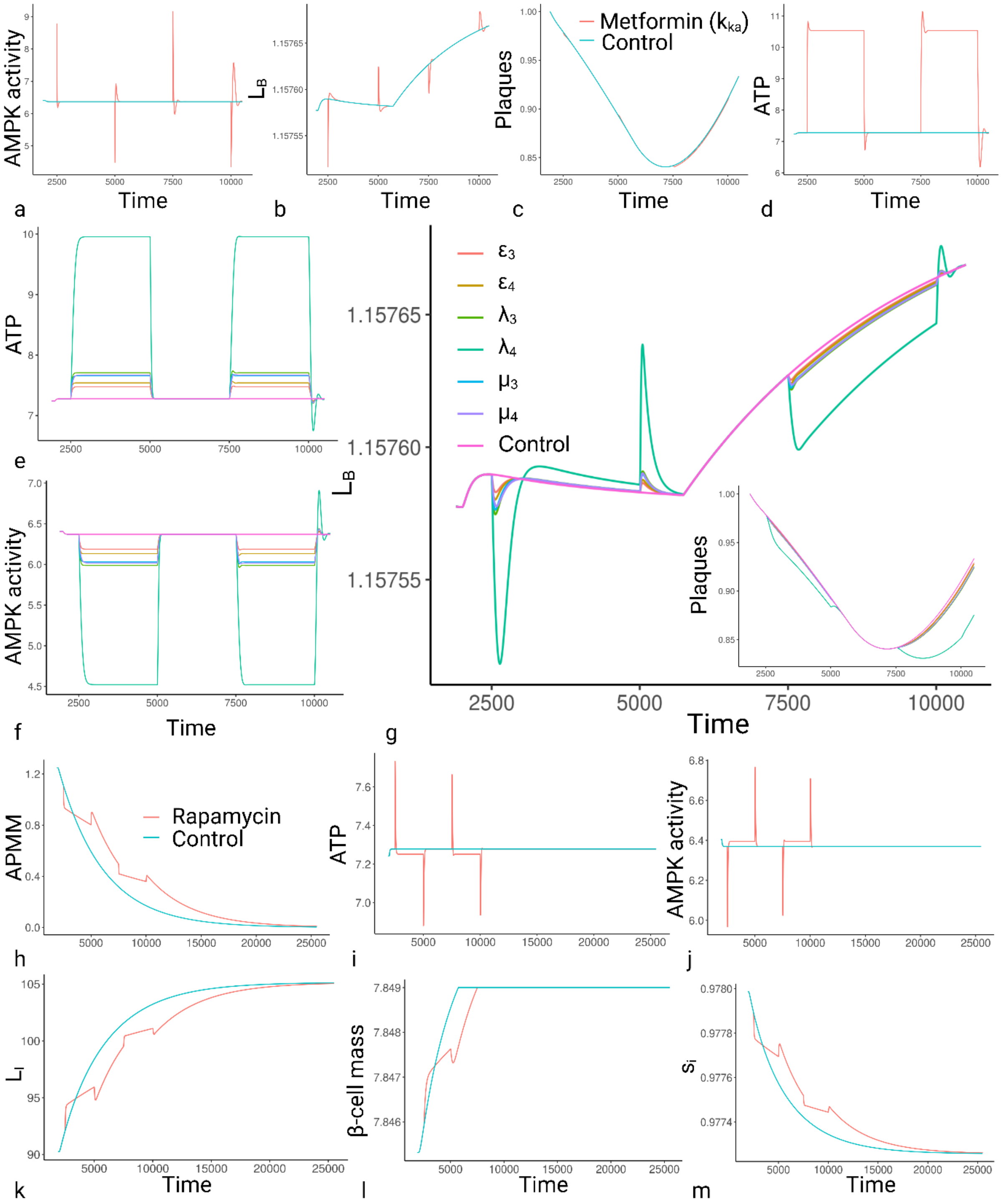


*Figure S5 | Effects of anti-ageing interventions on FEM variables and processes. (A-M) Treatments were two doses introduced between t = 2500 and t = 5000 au and then between t = 7500 and t = 10000 au. (A-D) Metformin modelled as a change in $k_{ka}$ for AMPK activation (or $k_{kd}$ for deactivation).* (A) AMPK activity. (B) $L_B$. (C) Plaques. (D) ATP. *(E-G) Metformin modelled as a change in parameters affecting AMPK regulation of IMC. (E) ATP. (F) AMPK activity. (G) $L_B$ and plaques (bottom right).* (H-M) Effects of rapamycin. *(H) APMM. (I) ATP. (J) AMPK activity. (K) $L_I$. (L) β-cell mass. (M) $s_i$. Abbreviations: AHCM, ATP hydrolysing cellular metabolism; APMM, ATP producing mitochondrial metabolism; ATP, adenosine triphosphate; FEM, Fuel and energy model; IMC, intrinsic mitochondrial capacity; $L_B$, blood lipid; $L_I$, intracellular lipid; $s_i$, insulin sensitivity.*

Increasing the rate of AMPK activation (or decreasing rate of deactivation) had little impact on the FEM. Most variables and processes spiked (up or down) at the point $k_{ka}$ (and $k_{kd}$) changed, but quickly returned to control levels, such as $L_B$ (Figure S5B), while the number and size of associated plaques showed little change (Figure S5C), despite shifting the equilibrium level of ATP (Figure S5D). Adding a variable for ATP-independent AMPK activation, $k_{in}$ (value 0 in control and 0.5 for metformin), as in equation S9, had the same lack of effect (data not shown).

$$\frac{dK}{dt} = k_{ka}\frac{AMP}{ATP} - k_{kd} \cdot K \ \rightarrow k_{in} + k_{ka}\frac{AMP}{ATP} - k_{kd} \cdot K (S9)$$

The results reflect that AMPK activity is not primarily affected by the rates of activation and deactivation (dependent or independent of ATP). AMPK activity is determined by the ratio of ATP:AMP, activating and deactivating when levels are out of equilibrium and returning to basal levels when equilibrium returns.

We therefore modelled changes in the parameters $\mu_3$ and $\mu_4$, $\lambda_3$ and $\lambda_4$, and $\varepsilon_3$ and $\varepsilon_4$, determining how AMPK regulates IMC. We multiplied or divided all parameter values by 1.5 so that each change either increased the AMPK-activated IMC or prevented the inhibition, as shown in Table S3. Results for ATP, AMPK activity, $L_B$, and plaques are shown in Figure S5E-G.

*Table S3 |IMC parameters and value changes representing metformin.*

| Parameter (code version) | FEM | FEM + Metformin |
|---|---|---|
| $\mu_3$ (mu3) | 0.021 | 0.0315 |
| $\mu_4$ (mu4) | 0.025 | 0.01667 |
| $\lambda_3$ (lam3) | 10.892 | 7.26133 |
| $\lambda_4$ (lam4) | 5.6 | 3.7333 |
| $\varepsilon_3$ (eta3) | 1.7 | 1.1333 |
| $\varepsilon_4$ (eta4) | 8.5 | 12.75 |

*Abbreviations: FEM, Fuel and energy model; IMC, intrinsic mitochondrial capacity.*

Thus, as all parameters for the impact of K on IMC produced similar curves, and $\lambda_4$ produced the greatest effect, we modelled metformin in Figure 4C-E using $\lambda_4$. As indicated in Figure S5G, when metformin was modelled as boosting AMPK-activated IMC, it produced only transient effect on $L_B$, but produced extended reduction of risk from plaque formation (as modelled by equation S3.10). The results suggest that if metformin acts by the mechanisms suggested here that it is not gero-protective, but risk preventative.

##### The impact of rapamycin on ageing as modelled by the FEM

We modelled rapamycin as discussed in the main article, as both an initial drop ($0.15 \cdot k_{CM}$) in $k_{CM}$, but also a reduction in the rate of $k_{CM}$ decline (0.25 times). The results are shown in Figure 5G-I, with additional variables and processes shown in Figure S5H-M. The results demonstrate that rapamycin induces initial drops in AHCM (Figure 5G) and APMM (Figure S5H) and rises in $L_I$ (Figure S5K) and basal inflammation (Figure 5I), consistent with a worsening metabolic profile and metabolic slowdown, but due to the decreasing rate of selective destruction these same processes and metabolites are improved in the longer term.

#### Model discussion

All models are simplifications of reality, and biological models in particular cannot hope to capture the full complexity of the systems they represent. However, here we have tried to demonstrate that homeostatic systems designed to maintain key metabolites at equilibrium will necessarily have knock-on effects leading to increases in either metabolites such as $L_I$ or processes such as inflammation. Notably, we have not calibrated the model against any specific biological systems, instead validating the model where possible by its consistency with age-related outcomes and interventions. Further work should include adding organism-specific detail to the model so that it can be properly calibrated to specific systems, but even without these additional steps the FEM clearly indicates that a network designed to maintain equilibrium ATP will have consequences for multiple aspects of fuel and energy homeostasis in response to metabolic slowdown.

We suggest that based on the biological evidence available, metabolic slowdown will lead to increasing $L_I$, $L_B$, basal inflammation, insulin resistance, and mitochondrial dysfunction, likely reflecting the underlying cause of age-related dysfunction and disease. However, the model is not without limitations or areas for expansion, which will be discussed below.

#### Metabolite equilibria and dynamic compensation

In the FEM, $L_B$, ACoA, and ATP are all maintained at equilibrium despite changes in multiple parameters and variables (see Figure S1). Changing meal and $k_{CM}$ caused spikes followed by swift return to equilibrium (as long as fuel remained in the adipose). This robustness to changing parameters was termed dynamical compensation by Karin, et al. (2016)[13], who modelled blood glucose. They observed that in the β-cell, insulin, glucose (BIG) model first described by Topp, et al. (2000)[24], changes in blood glucose caused corresponding changes in βM, via alterations in the rates of apoptosis and proliferation, which modulated the level of insulin so it could return blood glucose precisely to equilibrium.

As described in the main article and supplementary model construction, $L_B$ is regulated by insulin in a similar way to blood glucose, so we replicated the βM equation[13] for $L_B$ (S3.4). We used the same equation for IMC (S1.9) and its regulation by AMPK, which helped create the robustness of the FEM to maintain $L_B$ and ATP at equilibrium despite changes in meal values or $k_{CM}$ (and most other parameters, as shown in Figure S1). Only the μ, λ, and ε parameters influencing βM and IMC influence the equilibria for $L_B$ and ATP respectively.

However, in both cases, the equations S1.9 and S3.4 are unlikely to reflect the real dynamics as accurately as the equation for βM governing glucose in the BIG model. Glucose is a single, soluble molecule which is carried free in the blood, FFAs include multiple molecules and are not soluble in water, so only a tiny percentage (<0.01%) are carried free in the plasma, with most stored as esters transported on lipoproteins. Thus, while changes in blood glucose are a result of additional molecules entering or leaving the blood and must be counteracted by the opposite change, FFAs can be replenished from sources within the blood such as lipoprotein-bound triglycerides, and different FFAs may be regulated in different ways.

Importantly, the levels of lipoproteins such as LDL also increase with age, as do serum triglycerides, cholesterol, and various markers of $L_B$[66,67], suggesting that it does not have dynamic compensation, at least to the same level as blood glucose which only shows significant changes in the event of diabetes[24,68]. Thus, the regulation of $L_B$ is perhaps more similar to the regulation of $L_I$, where negative feedback shifts the equilibrium rather than maintains it (Figure S2E and S).

However, there is also good evidence that at the fundamental level described in the FEM, glucose and $L_B$ are regulated in a similar way: glucose increases β-cell survival[69] and proliferation[70], as do FFAs at least as far as proliferation[27-29], although the effect may be more complicated[71], and some FFAs can inhibit glucose-induced insulin secretion[72]. Whether FFAs protect against β-cell apoptosis is more controversial, with several studies suggesting that FFAs induce β-cell apoptosis[73,74]; however, this (lipo)toxicity effect is also observed for glucose (glucotoxicity), with low levels of glucose protecting β-cells from apoptosis while high levels induce apoptosis[75].

Notably, both glucose and FFAs have been implicated in diabetes and it is also clear that much of the inhibitory effects of FFAs on insulin production reflect prolonged exposure under diabetogenic and obesogenic conditions[74,76]. It is likely that low levels of FFAs protect β-cells from apoptosis, but again it may be complicated by the various types of FFAs. The dynamic compensation for $L_B$ may therefore be reduced or removed by factors outside the FEM, or LB may have different effects on insulin production to blood glucose even at the fundamental level, but the question remains of the relevance of dynamic compensation for the conclusions drawn here. We therefore modelled the FEM with constant βM (at different values), as shown in equation S10 and Figure S6.

$$\frac{d[\beta M]}{dt} = \beta M \left( \frac{\mu_1}{1 + (\frac{\lambda_1}{L_B})^{\varepsilon_1}} - \frac{\mu_2}{1 + (\frac{L_B}{\lambda_2})^{\varepsilon_2}} \right) \to 0 \quad (S10)$$

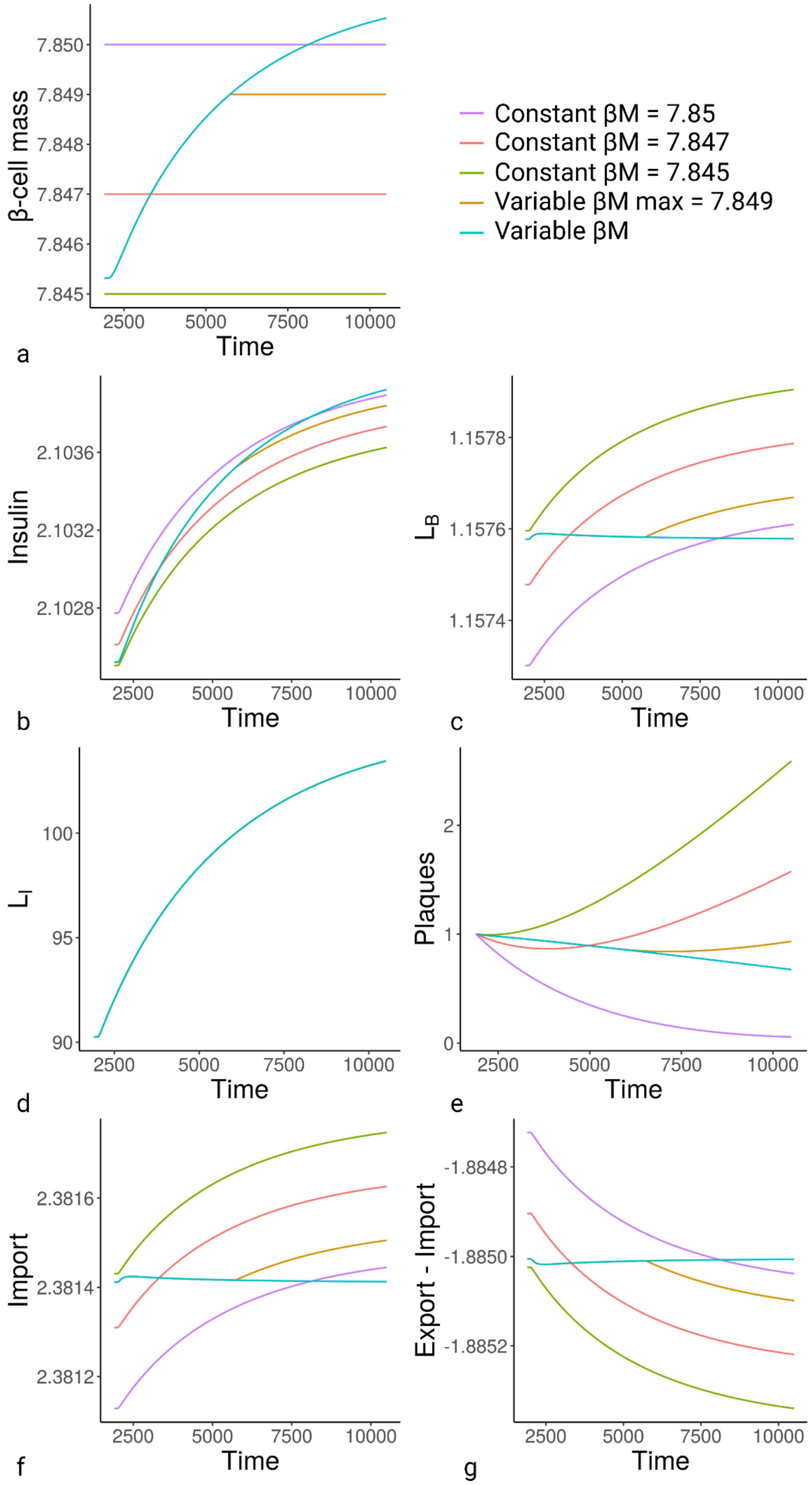


*Figure S6 |Impact of constant βM on FEM. (A) βM input. (B) Insulin. (C) $L_B$. (D) $L_I$. (E) Plaques. (F) Import. (G) Export minus import. Abbreviations: βM, β-cell mass; FEM, Fat & Energy Model; $L_B$, blood lipid; $L_I$, intracelluler lipid.*

Regardless of the starting value of βM, only variable levels allowed dynamic compensation for $L_B$ (Figure S6C). However, losing dynamic compensation for $L_B$ only accelerated the outcomes predicted by the FEM in response to metabolic slowdown, particularly the rise of $L_B$ (Figure S6C) and formation of atherosclerotic plaques (Figure S6E), without affecting $L_I$ or basal inflammation (Figure S6D and data not shown). **Thus, the ageing phenotype produced by the FEM is not likely to be a peculiarity of a particular model, but a robust effect of the co-regulation of fuel and energy pathways in response to the continuing unidirectional change of metabolic slowdown**.

Atherosclerosis does appear to occur in young obese individuals[77], but this may still reflect factors outside the FEM prohibiting the dynamic compensation. We maintained the dynamic compensation for $L_B$ in the FEM because it appears to be regulated by the pancreas and insulin similarly to blood glucose, which should produce the same dynamic compensation.

Notably, we could also have achieved a similar dynamic compensation for $L_I$ by modelling inflammation as a product of immune cells and TNF-α using similar equations as βM and insulin. However, there is no evidence for dynamic compensation for $L_I$. The equilibrium could only refer to the maximum $L_I$, as negative energy balance must always lead to $L_I$ depletion to provide fuel for energy. As mentioned in the main article, there is little evidence that the body increases energy expenditure through exercise (see below for heat) to compensate for overfeeding[78], or for the 'set-point hypothesis' that maximum weight is controlled neurologically by the hypothalamus[79], although it has been suggested that the set-point might be camouflaged by Western diets[64], specifically refined sugar[80]. Likely, the truth is more complex, reflecting increased fuel burning as $L_I$ increases without a set equilibrium, but this additional burning is blunted by Western diets.

Studies examining overfeeding indicate with equal clarity that one excess calorie absorbed does not equal one excess calorie stored[81,82]. In a recent systematic review of 19 studies, Bray and Bouchard (2020)[82] indicated an essentially linear relationship between excess calories in and additional energy stored (energy gain = 3,731 + 0.56 · overfed kcal). At the lower end of excess calories (~10,000 kcal), 1 excess kcal = 0.93 kcal stored, whereas at higher excesses (~80,000 kcal), 1 excess kcal = 0.61 kcal stored. While $L_I$ is clearly not regulated with the same stringency as ATP and $L_B$, these results indicate there are mechanisms to restrain $L_I$ growth during positive energy balance, and these mechanisms become increasingly active the higher the excess energy. Notably, it is unlikely to reflect a simple storage incapacity, as extra $L_I$ is stored at all intake values.

Another mechanism which could regulate $L_I$ occurs in brown adipose tissue (BAT), where mitochondrial uncoupling allows the futile cycling of electrons across the mitochondrial membrane, burning FFAs to produce heat without converting ADP to ATP. While interesting, the main purpose of uncoupling appears to be heat generation rather than burning excess calories, as it rises during cold stimulation dependent on the presence of BAT[83], was higher under ambiently colder conditions[84], and most importantly was blunted by obesity[85], which is the opposite to what you would expect if it was a major pathway for the removal of excess fat. Equally, clinical studies examining why excess calories led to smaller gains in energy and fat mass than expected (metabolic inefficiency), found little evidence for an increase in basal metabolic rate (or physical activity) expected by an increase in thermogenesis[86]. Although others suggest "thermogenic activity offers tremendous potential to combat obesity"[87], increasing in response to high calorie diets to provide obesity resistance[88,89].

While these findings are significant in the combat of obesity, they have little impact on the conclusions drawn from the FEM, as they can be factored in as part of the equation introduced by Bray and Bouchard (2020)[82], reducing the additional weight gain as excess calories increase. Uncoupling may reduce the rate at which $L_I$ increases in response to metabolic slowdown, but it does not prevent it. The (more slowly) increasing $L_I$ therefore still induces hypertrophy, IR, basal inflammation, and the eventual increases in $L_B$ once lipid storage becomes exhausted. As brown adipocytes have fixed cell number, this will also limit their capacity for $L_I$ removal. Indeed, any system, such as lipid removal in the bile (RCT), that cannot be increased inexhaustibly in response to continuing unidirectional changes like metabolic slowdown, will eventually hit a maximum that prevents further restoration of equilibrium. The difference in outcome is if equilibrium is maintained until the threshold is exceeded, it will result in a sudden change in metabolite dynamics as we have modelled for $L_B$,and risk from plaques, or if the metabolite is allowed to continually change over time as we modelled for $L_I$ and basal inflammation. Speculatively, the former may reflect the induction of age-related disease (as stricter homeostasis likely corresponds to increasingly important metabolites), and the latter simple age-related decline.

Contrary to $L_I$ and $L_B$, there is good evidence for a robust ATP equilibrium that does not change with age, as discussed in the model construction and main article. In the FEM, this is maintained by the regulation of IMC by AMPK as in equation S1.9. As AMPK is a function of ATP:AMP ratio, this maintains equilibrium ATP via the same mechanism as the βM equation S3.4 does for $L_B$: when ATP decreases, IMC increases to elevate ATP production and restore ATP; and when ATP increases, the IMC decreases reducing ATP production and restoring ATP. However, it should be noted that while the ATP levels remain constant, the size of the adenosine phosphate (AP) pool increases with age[90], suggesting that our equations S1.5-1.7 are oversimplifications.

The main oversimplification for ATP dynamics is grouping all the factors affecting intrinsic mitochondrial capacity together into a single variable, intrinsic mitochondrial capacity (IMC), when the literature suggests it is affected by multiple processes and regulators, including several that are thought to be integral to the ageing process, such as autophagy. Further research into how exactly the IMC is being reduced could have significant utility in the development of anti-ageing therapies. Reversing the decrease in IMC could prevent the shift in ACoA dynamics that leads to insulin resistance, weight gain, basal inflammation, and atherosclerosis.

In the FEM, ACoA is kept at equilibrium (Figure 4D) despite the absence of an equation specifically designed to induce dynamic compensation. As we observed when implementing equation S8 for β-oxidation, which removed IMC (in ACoA formation), the dynamic compensation results from ACoA *formation* being controlled by the same IMC inducing its *deacetylation* (i.e. degradation) during APMM. As mentioned above, the complexities of IMC could potentially separate the intrinsic capacity for β-oxidation from the intrinsic capacity for APMM. Further research is required to address this. However, as we showed in Figure S4B-F, whether or not ACoA remains at equilibrium in the event of metabolic slowdown is immaterial to the conclusions of the FEM. A rise in ACoA in response to declining AHCM will still result in increased FFA synthesis, insulin resistance, weight gain and atherosclerosis. As we show repeatedly in this analysis, changes to the model equations affecting the specifics of metabolite and process regulation do not prevent the inevitable shifts in homeostasis from metabolic slowdown.

#### Regulation of $L_I$

A key function of the adipose tissue is exporting lipid back into the blood during negative energy balance, allowing non-adipose tissues to maintain ATP production via β-oxidation. In equation S3.5, export is inhibited by insulin, but in reality there are many additional regulating factors. Adipose tissue is one of the few tissues along with the liver that responds to glucagon, stimulating lipolysis in rats[91] and the release of FFAs and glycerol into the blood[92] in a mechanism that is inhibited by insulin[93,94]. However, the relationship is likely more complicated in human adipose than rodents[95], and some studies indicate glucagon may actually reduce blood lipid[96,97]. Clinical studies showed no effect of glucagon on adipose lipolysis in people with and without diabetes[98], and expression of white adipose tissue (WAT) glucagon receptor did not affect lipolysis or hepatic lipid accumulation in mice[99]. There is also evidence for roles of somatostatin and adrenaline in regulating lipolysis and export, but due to the complexities of these regulatory systems, and the likelihood that they regulated export in response to conditions outside the FEM, such as stress, we chose not to include them. However, understanding the significance of glucagon and other hormones in adipose homeostasis may help ameliorate the impact of metabolic slowdown on lipid accumulation.

Another simplification of the FEM regarding $L_I$ was described in equations S3.6-3.8. As we reduced the model to include only the adipose and the blood, the model did not respond to metabolic slowdown with compensatory changes to import and export (Figure S2L-N) as it did to changes in meal value (Figure S2Z-iii). Metabolic slowdown affected only the adipose rate of lipid β-oxidation without impacting additional tissues. The latter would have altered rates of non-adipose import, necessitating compensatory changes in adipose import and export to maintain $L_B$. Further development of the model should therefore include a multi-tissue system.

Probably the least realistic simplification employed in the FEM is the simulation of inflammation as a single cytokine, TNF-α, which is produced only by hypertrophic adipocytes. It would be impossible to include a realistic model of the immune system without it dominating all other components. However, further work should include factors such as hypertrophic adipocytes secreting monocyte chemoattractant protein-1 (MCP-1) and granulocyte colony stimulating factor (G-CSF)[100], both of which attract immune cells, and the latter mobilizes haematopoietic stem cells for the production and release of neutrophils from the bone marrow[101,102], expanding the pool of monocytes and macrophages, and enhances their phagocytic function[103,104]. The incorporation of immune cells and their relationship with adipocytes would provide further understanding of the role of inflammation in adipose homeostasis, but unlike many other studies which interpret inflammation as largely destructive or of unknown function, the FEM suggests a clear function for inflammation in removal of excess fat, insulin sensitization, and the resultant maintenance of lipid homeostasis.

The danger with any model of inflammation is that it spirals out of control, creating a vicious cycle that is purely destructive. There are perhaps many in the field of gerontology who have this view of inflammation, but in reality inflammation has a series of robust negative feedback mechanisms that largely prevent these vicious cycles. The FEM suggests that rising basal inflammation reflects a mixture of mitochondrial dysfunction and lipid dysfunction, and need not result from damage accumulation or allostatic load[105,106]. However, these causes are not exclusive with each other.

One seemingly vicious cycle we did not include in the FEM is the link between inflammation and insulin resistance, which is so strong that there is great uncertainty as to which is cause and which is effect. A role for IR in inducing inflammation is consistent with inflammation as a mechanism of removal of insulin resistant hypertrophic cells as predicted by the FEM. Thus, evidence such as the inhibition of mTORC2 and resultant IR inducing increased polarisation to inflammatory M1 macrophages through MCP-1[107] is not surprising. However, that inflammation induces IR is counterintuitive to the mechanisms proposed here.

Macrophage infiltration and activation has been shown to induce insulin resistance in adipose tissue and in vitro adipocytes[108,109]. Equally, knockout of IκB kinase-beta (IKK-β) or c-Jun N-terminal kinase 1 (JNK1) involved in macrophage signalling protected mice from diet-induced insulin resistance[110-112]. Importantly, in many of these studies, genetic changes in the macrophages were capable of inducing

IR in adipocytes, suggesting the IR was not just a byproduct of obesity which also induced inflammation; the inflammation caused the IR.

It should be noted that many of the conditions demonstrating inflammation induces IR were heavily biased against the insulin-sensitizing effects of inflammation, such as obesity or in vitro conditions with high numbers of immune cells. It is likely that if the macrophages were in a healthy, non-obese tissue where resolution of the excess $L_I$ could occur, they would deactivate and exfiltrate, promoting the anti-inflammatory polarisation we have discussed above where dead adipocytes are replaced by new insulin sensitive ones. However, in obesity and in vitro, resolution can be prevented by multiple conditions including immune cells at supraphysiological levels or constitutive activity due to mutation, or the absence of the stem cells to replace insulin resistant adipocytes with new insulin sensitive ones. Under such conditions several factors may raise the level of immune activation so that it is more resembling of an immune response to infection rather than basal inflammation to remove hypertrophic cells.

Crucially, just as the body benefits from reducing ATP production in infected cells, by prohibiting pathogen replication, it also benefits from reducing import of fuel into infected cells, which could instead be absorbed by the immune cells combatting the infection, rather than feeding pathogen infected cells that may aid pathogenic replication. It therefore makes evolutionary sense that higher levels of inflammation consistent with infection – which may occur in obese individuals where the mounting immune response is insufficient to clear the excess $L_I$ (and thus deactivate) – induce IR.

In conclusion, the FEM makes multiple predictions that will require validation both in the laboratory and with more detailed modelling to include the aspects discussed here. However, many of the metabolic changes predicted by the FEM are not dependent on specific equations or parameter values, and include only variables well described in the biological literature. They strongly predict that if AHCM declines with age, the resulting metabolic cascade is unavoidable and will result in dysfunction, disease, and death if not prevented.

#### Expected Outcomes and Falsifiability

The predicted outcomes of future experiments and interventions according to SDT, as well as outcomes which would falsify it are presented in Table S4. However, it should be noted that the expected outcomes are not necessarily exclusive to other theories, and further work is required, including more detailed models of damage accumulation, to elucidate which outcomes are predicted by which theory.

*Table S4 | Expected and falsifying outcomes for SDT from future experiments and interventions*

| Selective destruction results in metabolic slowdown | |
|---|---|
| Expected Outcomes | |
| 1. | Mechanisms by which cells can determine the growth rate of the surrounding cells and compare this to their own will be discovered. |
| 2. | These mechanisms will be connected to pathways regulating epigenetics, thus allowing one cell to influence the epigenetics of its neighbours. |
| 3. | Slower cells will be more likely to influence the epigenetic landscape of faster cells or have a stronger effect than that of faster cells on the epigenetics of slower cells. |
| 4. | The epigenetic changes should be causative in metabolic slowdown, and occur at the loci which correlate with biological age in the epigenetic clocks. |

| | |
|---|---|
| 5. | There should not be significant difference between an aged cell and a cell whose metabolism has been slowed epigenetically, except for telomere length and stochastic changes which promote noise. |
| Falsification | |
| 1. | Observations from independent labs that, in conditions including in vivo, slower cells (including stem cells) either accelerate or have no effect on the metabolism and/or epigenetic landscape of faster cells. |
| 2. | Observations from independent labs that removing the epigenetic markers of aged cells in animals is associated with no additional mortality related to overactivity disorders and/or reduces the rate of AHCM. |
| 3. | Discovery of a mutant control system in cells and tissues which is sufficient to control all fast mutants (perhaps autonomously or by other mechanism) without providing a further fitness benefit to the organism by giving slower cells a competitive advantage. |
| Metabolic slowdown induces ageing and age-related disease | |
| Expected Outcomes | |
| 1. | Interventions reducing AHCM will not affect ATP levels in the medium and long term because ATP production will also decline. |
| 2. | The mechanisms reducing ATP production in ageing precipitate mitochondrial dysfunction. |
| 3. | Reduced number of adipocytes, but not necessarily removal of adipocytes (which could result in adipocyte proliferation and return to homeostasis), should result in a lower capacity to store $L_I$, leading to increased IR and higher risk of atherosclerosis. |
| 4. | Mechanisms removing excess lipid from the blood will be important in ageing and age-related diseases in multiple organisms. The propensity of individuals for these pathways, such as RCT in the liver or uncoupling in the BAT, as well as lipid transport within the blood, will correlate with risk of developing various diseases of ageing, particularly atherosclerosis. |
| | NB: As SDT would predict, low LDL cholesterol is associated with lower atherosclerotic risk, but higher cancer risk[113,114]. Mendelian randomisation showed low LDL was unlikely a causal factor in cancer[115], suggesting it is underlying factors which promote both low LDL and cancer independently. One such factor could be the rate of selective destruction. |
| 5. | Reducing or inhibiting basal inflammation in non-obesogenic conditions should have negative effects on adipose homeostasis, including increased hypertrophy and IR. |
| 6. | The impact of dietary changes on $L_I$ and $L_B$ will be complex due to compensatory homeostatic mechanisms, but reducing lipid or even total calorie intake would be expected to have little impact on $L_B$ in younger individuals before the maximum thresholds of βM/RCT/uncoupling are reached, but more of an effect in older individuals with slower metabolism and more difficulty upregulating the compensatory mechanisms. |
| Falsifiability | |
| 1. | Evidence from independent labs in longitudinal studies that ATP levels drop or AHCM increases with age in multiple tissues of species and/or mutants which show otherwise normal ageing (including IR, basal inflammation, and age-related disease). |
| | NB: Evidence of additional systems (such as RCT) which can buffer the changes in homeostasis described here, or evidence that metabolites modelled here to be in homeostasis instead change, with age should not falsify the conclusions of SDT. Falsification by this mechanism would require demonstration that regulation of ATP, ACoA, and lipid metabolism are independent. |
| 2. | Each point along the chain of causation could be falsified by studies indicating alternative mechanisms of regulation which do not require homeostatic shifts associated with ageing and age-related disease.<br>For example:<br>• If some species or mutants utilise uncoupling to maintain $L_I$ without immune involvement, but still have escalating basal inflammation with age, this would suggest |

| | |
|---|---|
| | that some process other than immune regulation of adipose is the primary cause of basal inflammation.<br>• If some species or mutants show levels of $L_B$ that do not increase with age, without major changes in the types of lipoprotein carriers or specific lipid profiles, but still show increasing plaque formation, then this would suggest that some other factor, such as endothelial dysfunction, is the primary cause of atherosclerosis. |
